# A mismatch in wildfire impact reduction and ecological restoration in western U.S. fuel treatments

**DOI:** 10.64898/2026.10.30.763129

**Authors:** Timo Metz, Nicholas Hendershot, Chuliang Song

## Abstract

A century of fire suppression has made historically fire-adapted forests across the western U.S. denser and dominated by fire-intolerant trees, leading to severe fires. Fuel treatments aim to reduce wildfire impact and restore fire-adapted forests. Ecological theory suggests these objectives need not align: protecting the existing, fire-maladapted stand may prevent fire-tolerant seedling recruitment. Using Forest Inventory and Analysis data on 81,621 trees and 41,328 seedlings across 11 western U.S. states, we show that fuel treatments increase tree survival, but benefits are largest for fire-intolerant trees, fade within nine years, and come with no seedling composition shift towards fire-tolerant species. Concluding, fuel reduction reduces wildfire impacts but does not restore fire-adapted forests. Without changes in treatment design, long-term resilience will require repeated intervention.

## Introduction

Wildfires have drastically increased in past decades across the western United States, threatening livelihoods, biodiversity, critical water resources and the long-term persistence of forests on the landscape (*1–3*). Climate change further intensifies these challenges, driving more severe wildfires and prolonged droughts that risk the conversion from forest to shrub- or grassland after fire (*3–6*). The increased risk of high-severity, stand-replacing fires across much of the western U.S. is driven by decades of fire suppression and logging practices which have made historically frequent-fire forests more dense, homogeneous and dominated by fire-intolerant species with high amounts of accumulated fuel (*7–9*). In response, federal and state agencies invest billions of dollars annually in forest management strategies such as mechanical fuels reduction treatments and prescribed burns to decrease fuel loads and thereby lower wildfire severity (*10*, *11*). Recent studies have shown that treatments effectively reduce fire severity and associated damages (*12–14*). However, treatments also have a second stated goal: restoring fire-adapted forest composition and structure, so that forests can withstand recurrent wildfire in the long-term without repeated intervention (*10*, *15–17*). These two objectives of treatments — fire hazard reduction and forest restoration — are usually viewed as complementary, because reducing fire severity in the short term is assumed to promote resilience in the long term (*18*).

However, ecological theory suggests that the two objectives of fuel treatments need not align. Resilience, which is the capacity to maintain state and function in the face of disturbances, can be decomposed into two components: resistance (the ability to withstand disturbance) and recovery (the capacity to return to pre-disturbance levels) (*19*, *20*). In historically frequent fire forests, such as ponderosa pine or Sierra mixed conifer forests, recurrent low- and moderate-severity fires maintained a feedback in which each fire reduced surface fuels, preferentially killed fire-intolerant trees and opened canopy gaps in which seeds of fire-tolerant trees, which are often shade-intolerant, could germinate (*21*, *22*). Resistance to fire and recovery towards a fire-adapted composition thus reinforced each other, maintaining resilience in the long term. In infrequent stand-replacing fire forest types that are naturally dense and dominated by fire-intolerant species such as lodgepole pine, fir or aspen, resistance is low, but intervals between fires were historically long enough to allow recovery (*23*, *24*).

Fuel treatments effectively copy the fuel-reducing role of fire (*11*), but we hypothesize that in historically frequent-fire forest types they may fail to restore a fire-adapted forest composition. That is because treatments aim to preserve the existing stand, which in fire-suppressed forests is dominated by fire-intolerant trees, leading to seedling recruitment likely mirroring the preserved overstory rather than shifting towards fire-tolerant species. As fuels rebuild (*25*), the forest then quickly returns to the pre-disturbance conditions that made it vulnerable in the first place (Fig. 1a). This should be especially true for mechanical treatments that lack the element of primarily killing fire-intolerant trees unless directly targeting them for removal. However, they are the dominant form of fuel treatment across much of the western U.S. (*26*, *27*). Local studies are consistent with this proposed mechanism (*28–30*), but whether this is a general feature of contemporary fuel treatments across the western U.S. is unknown.

**Fig. 1:**
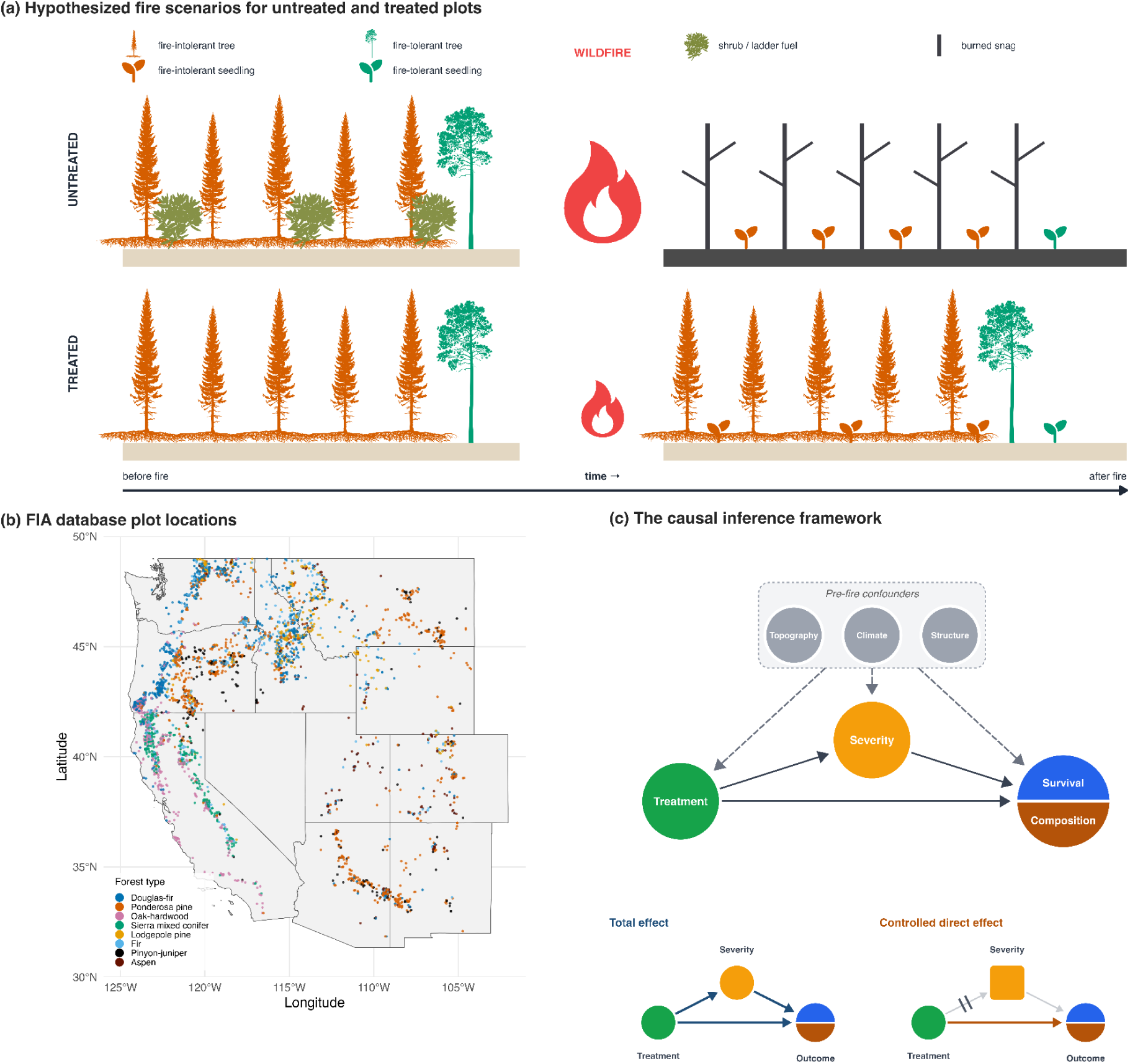
Mechanisms of treatments, distribution of FIA plots across western United States and investigated resilience mechanisms. (a) Untreated stands have developed high amounts of accumulated fuels and a dominance of fire-intolerant trees driving more severe, stand-replacing fires. Dominance of fire-intolerant trees in the landscape makes recruitment of fire-intolerant seedlings most likely. Treatments reduce accumulated fuels but preservation of the fire-maladapted overstory may also lead to fire-intolerant seedling recruitment and hence prevent a shift towards a fire-adapted forest. (b) Geographic distribution of plots used in this study. Locations are not exact. (c) The causal inference framework is used to estimate the total effect of treatments on tree survival and seedling composition shift and the controlled direct effect when holding fire severity constant. Pre-fire confounders (topography, climate and forest structure) are controlled for. Silhouettes are from PhyloPic (www.phylopic.org), flame icon from Bootstrap icons (www.icons.getbootstrap.com) and seedling icons from Font Awesome (www.fontawesome.com).

Here, we test whether pre-fire mechanical fuel treatments increase long-term forest resilience not only by making the existing stand more resistant, but also by shifting post-fire regeneration toward fire-tolerant species. Our analysis is based on the Forest Inventory and Analysis (FIA) tree and seedling inventory across 11 western U.S. states (see Fig. 1b). The tree data comprise 81,621 trees on 3,629 burned plots, and the seedling data 41,328 seedlings on 1,761 burned plots, covering fires between 1990 and 2023. Because treatments are applied in both frequent- and infrequent-fire forests, the data allow us to contrast forests where lower severity and a shift toward fire-tolerant seedlings are goals with forests where neither is expected. Treatments in the FIA data are mechanical, reflecting their dominance across much of the U.S. We use a causal inference approach to separate the total effect of treatment, including its effect through reduced severity, from its effect at a fixed severity. This lets us ask whether treatments shift seedlings toward fire-tolerant species under the surface fire conditions they are designed to promote (see Fig. 1c). Because treatments are not placed at random, we compare treated and untreated burned plots after adjusting for confounders that influence both where treatments are placed and how fires behave (topography, climate, and forest structure). We further assess the longevity of treatment effects in terms of tree survival and seedling composition shift and test regional-scale differences in treatment effectiveness across climate gradients.

## Results

### Treatments increase tree survival directly after treatments, but benefits decay rapidly

Fuel treatments substantially increase forest resistance by enhancing tree survival and decreasing crown fire probability (see Fig. 2a). Treatments nearly triple the odds of tree survival (total effect OR: 2.78, 95% CrI: 1.42-5.45) and treated plots also have 53% lower odds of burning at crown severity (OR: 0.47, 95% CrI: 0.33-0.65). Holding fire severity fixed, the effect is smaller (controlled direct effect OR: 1.82, 95% CrI: 0.91–3.61 at surface fire; OR: 2.45, 95% CrI: 0.91–6.64 at crown fire; Fig. S3a). This shows that part of the benefit comes from lower fire severity, and part may come from other mechanisms, such as decreased competition for resources. Treatment-related benefits in terms of odds of survival are largest for species with low fire tolerance (pooled OR 2.64, 1.69 and 1.34 for low, moderate and high fire tolerance, Fig. 2b).

**Fig. 2:**
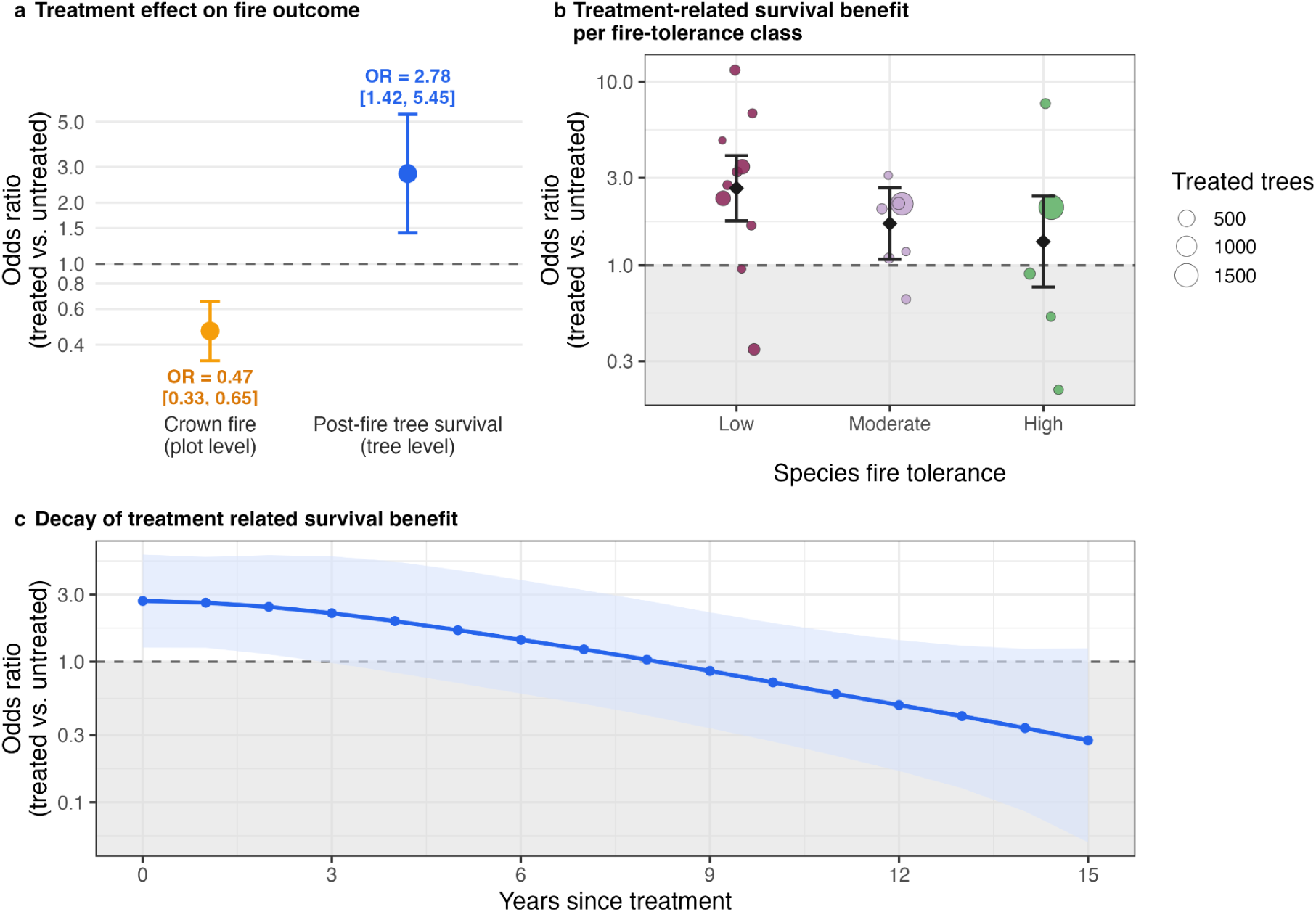
Treatment effects on crown fire probability and tree survival. (a) Total effect of treatments on crown-fire probability and tree survival. Points are posterior means, error bars are 95% credible intervals. (b) Controlled direct effect of treatments at surface fire for trees with low, moderate or high fire tolerance. Colored dots represent posterior means for each species, black diamonds are inverse-variance pooled ORs per class. Error bars on diamonds are 95% confidence intervals. (c) Decay of the controlled direct effect of treatments at surface fire over time, pooled over all forest types. The line is the posterior mean, dots mark years since treatment. The band in (c) represents the 95% credible interval. Grey area in (b) and (c) marks an OR < 1.

An assessment of treatment longevity shows that the survival benefit is short-lived (Fig. 2c). The benefit is large in years directly following treatment (OR = 2.74) but declines quickly, reaching OR = 1 (no benefit) between 8 to 9 years after treatment (see Fig. 2c). Beyond that point, the estimate falls below 1, indicating a potentially negative effect of treatments on survival, although the credible intervals include 1. How long the benefit lasts differs among forest types, from 5.7 years in Douglas-fir and pinyon-juniper to 13 years in ponderosa pine, with Sierra mixed conifer forests (7.4 years), aspen (7.5 years), lodgepole pine forests (9.4 years) and fir (9.5 years) in between, while oak-hardwood forests showed no detectable benefit at any time since treatment (Supplementary Fig. S1). Within a forest type, treatments help survival most on the driest plots (OR ≈ 6.1 on the driest, versus ≈ 0.8 on the wettest; OR 1.91 per global water-deficit SD, 95% CrI 1.03–3.55; Supplementary Fig. S2).

The total effect of treatments on tree survival is robust across estimators (Supplementary Fig. S3). To explain away the point estimate, an unmeasured confounder would need a risk-ratio association of about 2.7 with both treatment and survival, stronger than that of any measured confounder (Supplementary Table S1), and a negative-control outcome test on time since fire shows no detectable systematic difference between treated and untreated plots beyond what our models account for.

### Fire severity, but not fuel treatments, shifts seedling regeneration

Fuel treatments do not detectably change the share of seedlings in any fire-tolerance class. The total effect on the highly fire-tolerant fraction is +1.1 pp (−2.1 to +4.6), on the moderately tolerant fraction −2.5 pp (−7.4 to +2.2) and on the low fire-tolerance fraction +1.3 pp (−2.2 to +5.2) (Fig. 3a). All three posterior distributions are centred close to 0. Comparing plots that burned at the same severity gives the same answer. Under surface fire (the fire behaviour treatments aim to promote), treatment shifts no tolerance class by more than 1.2 pp. Under crown fire, the controlled direct effect is larger, with a decrease in the moderately tolerant fraction (−11.9 pp, 95% CrI: −21.8 to −0.4) that is redistributed to both the highly tolerant (+6.9 pp, −1.7 to +16.8) and the low tolerant fraction (+4.6 pp, −3.9 to +13.1). Apart from the moderate fraction under crown fire, all compositional shifts span 0 within credible intervals (see Fig. 3b). Fire severity itself, by contrast, has a relatively strong effect on composition in untreated plots. Untreated stands that burned as crown fire have 6.1 pp higher fraction of seedlings of species with high fire-tolerance (95% CrI: 3.8-8.6) and 5.1 pp fewer seedlings of species with moderate fire tolerance (95% CrI: −8.1 to −1.9) than stands burning at surface severity. The low fire-tolerance fraction is unchanged (−1.0 pp, 95% CrI: −3.5 to +1.3; Fig. 3c). Across plots measured 2-10 years after fire, predicted composition drifts in opposite directions in treated and untreated plots (Supplementary Fig. S4). In treated plots, the low-tolerance fraction falls from 37% to 28%, while the high-tolerance fraction rises from 7% to 10%. In untreated plots, both trends are weak and in the opposite direction, with the low tolerance fraction rising from 32% to 35% and the high-tolerance fraction declining from 8% to 7%. The moderate fraction rises from 53% to 59% in treated plots and declines slightly from 59% to 56% in untreated plots. However, the 95% credible intervals for treated and untreated plots overlap over the full period.

**Fig. 3:**
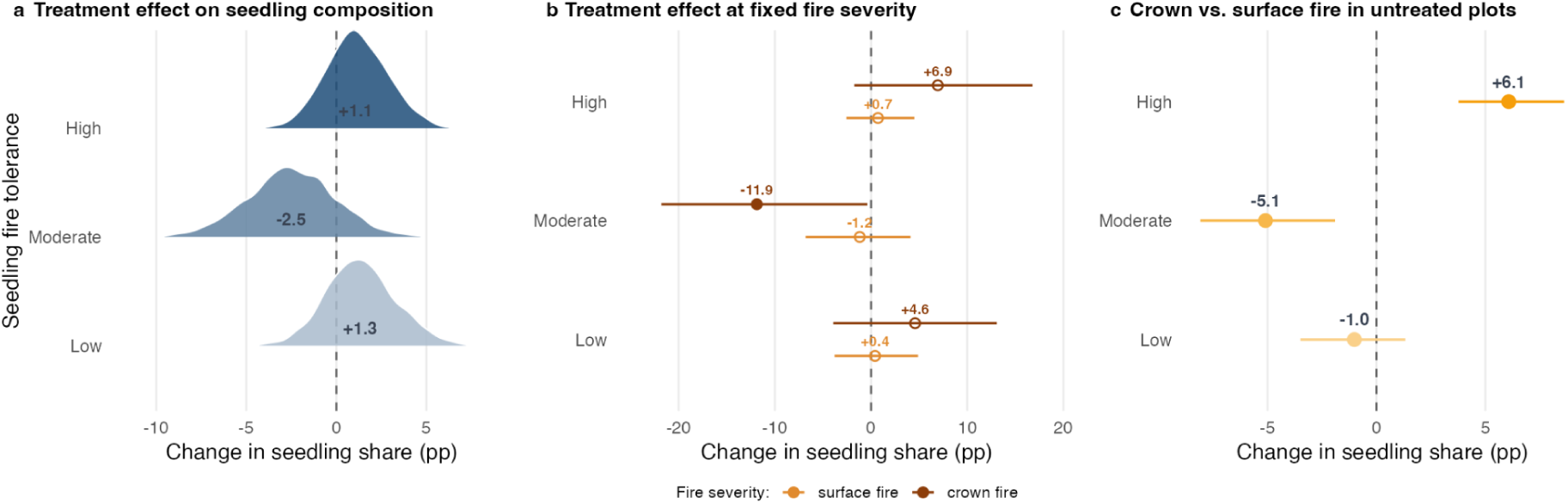
Fuel treatments do not shift post-fire seedling composition. (a) shows the total effect where densities are posterior distributions and labels posterior medians. (b) shows the controlled direct effect at surface and crown fire. (c) shows the seedling composition shift in crown compared to surface fire on untreated plots. The change in the share of seedlings per fire-tolerance class is based on marginal standardization, error bars indicate 95% credible intervals, both based on 2,000 joint posterior draws. n=1,761 plots (total 138 treated/ 1,623 untreated; crown fire 26 treated / 445 untreated) and 41,328 seedlings.

Comparing drier and wetter plots within a forest type, the treatment-induced shift towards fire tolerance is stronger on wetter compared to drier plots. The treatment effect on the highly fire-tolerant fraction goes from +2 pp on the wettest plots to roughly −18 pp on the driest plots, with the low and moderate fractions rising correspondingly. However, credible intervals overlap 0 along the whole gradient (Supplementary Fig. S5).

Across design-based estimators (Supplementary Fig. S6), the absence of a reduction in the low fire-tolerance fraction was robust, but the two reweighting estimators show a very modest gain of highly fire-tolerant seedlings and a reduction of moderately fire-tolerant seedlings. A negative-control outcome test on time since fire shows no detectable systematic difference between treated and untreated plots beyond what our models account for.

### Resistance and fire-adapted regeneration are mismatched in all studied forest types

The increase in tree survival without a substantial shift in seedling composition is consistent across forest types (Fig. 4). Treatment effects on tree survival are positive for Sierra mixed conifer, Douglas-fir, ponderosa pine, pinyon juniper, lodgepole pine, aspen and fir forest types. Oak hardwood is the only forest type in which treatments reduce tree survival. In ponderosa pine and lodgepole pine, the estimated seedling shift points towards fire tolerant species, but the 95% credible interval for the change in the dominance gap (fraction of seedlings with high fire-tolerance minus fraction of seedlings with low fire-tolerance in Fig. 4) does not exclude 0 (see faded vs. solid symbols in Fig. 4). In all other forest types there is no visible shift in the dominance gap. Treatment moves no forest type into a different quadrant of Fig. 4. Sierra mixed conifer and Douglas-fir, historically frequent-fire forest types where a shift would matter most, are dominated by fire-intolerant seedlings without treatment and remain so with it. Lodgepole pine, aspen and fir sit near the bottom of the y-axis with and without treatment, having a strong dominance of fire-intolerant species that is expected for forests with infrequent, stand-replacing fires. Dry ponderosa pine and pinyon-juniper are the only types dominated by fire-tolerant seedlings even without treatment, consistent with them being fire-adapted forest types.

**Fig. 4:**
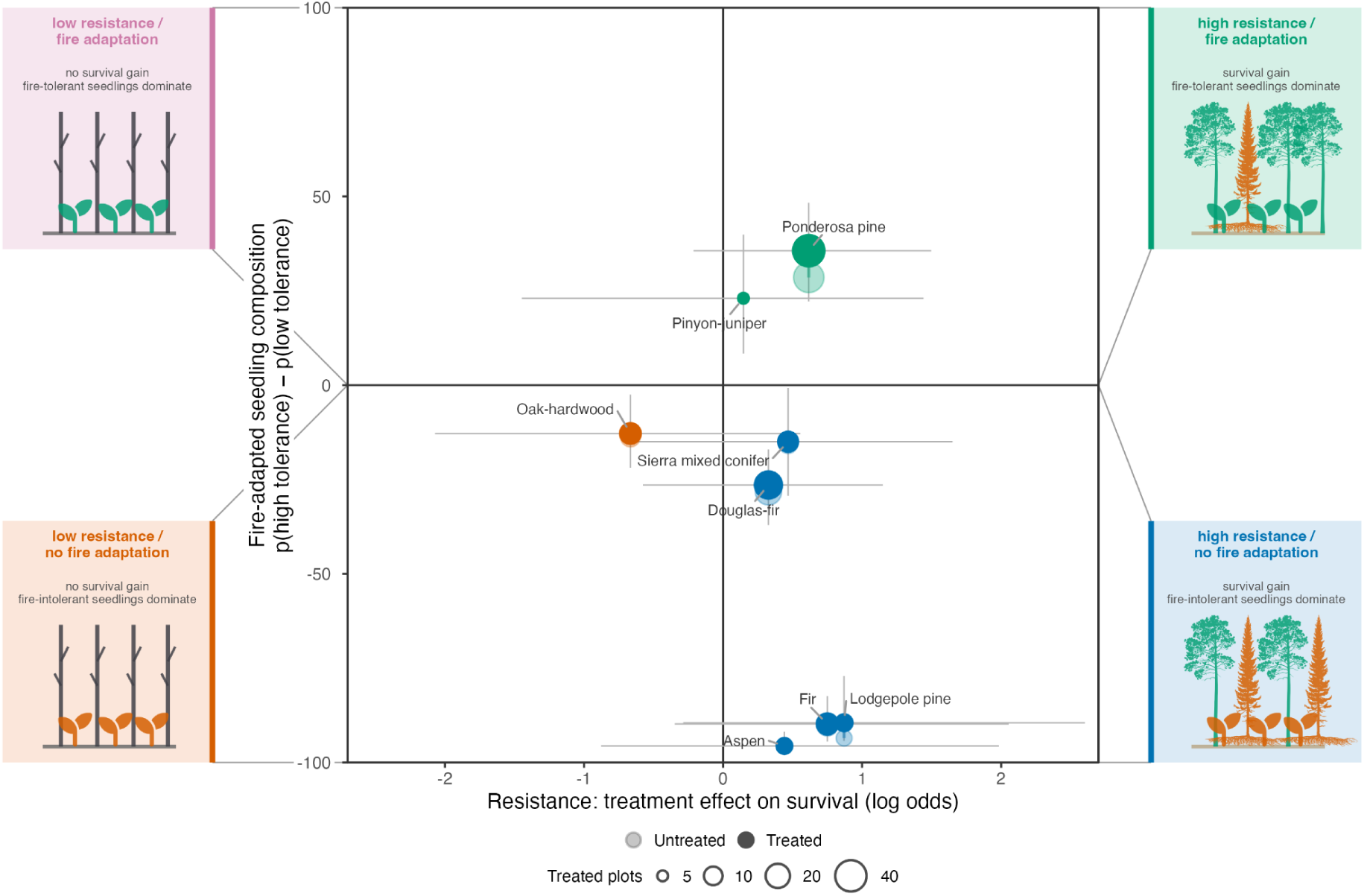
Resistance and fire-adapted regeneration across forest types. x-axis shows the controlled direct effect of treatment on tree survival at surface fire for each forest type. y-axis shows the predicted dominance gap at surface fire, i.e. expected fraction of a plot’s seedlings that belong to species with high fire tolerance p(high tolerance) minus the expected fraction of a plot’s seedlings that belong to species with low fire tolerance p(low tolerance). Error bars indicate 95% credible intervals. Colors indicate different quadrants. Quadrants indicate the four possible combinations of treatment effects in either increasing or decreasing tree survival (resistance) and dominance gap (fire-adapted regeneration). Faded dots show untreated plots and solid dots treated plots. If only a solid dot is shown, treated and untreated plots do not differ for that forest type. Silhouettes are from PhyloPic (www.phylopic.org), flame icon from Bootstrap icons (www.icons.getbootstrap.com) and seedling icons from Font Awesome (www.fontawesome.com).

## Discussion

Our findings show that across the western United States fuel treatments increase forest resistance to wildfire but do not restore fire-adapted regeneration. Treatments consistently increase tree survival and reduce crown-fire probability, but the survival benefit is largest for fire-intolerant trees and fades within nine years, well inside typical re-entry intervals (*12*, *25*). Treated plots, in other words, quickly rebound to their fire-maladapted state. Furthermore, we observe no shift towards fire-tolerant regeneration under the surface-fire conditions that treatments are designed to promote. Reducing fuels buys time, but by itself does not place forests on a trajectory towards naturally self-sustaining, fire-adapted communities. This pattern was previously documented only locally (*28–30*) but we show here that it holds across much of the western United States. Hence, our results suggest that protecting forests from severe wildfire in the short-term and restoring fire-adapted forests are related but distinct management objectives. Current fuel treatments effectively achieve the first, but by themselves do not appear sufficient to achieve the second.

This should not be interpreted as evidence that fuel treatments are counterproductive. Their primary objective is to reduce tree mortality during wildfire, and achieving that objective provides substantial benefits, particularly where the alternative is stand-replacing fire and potential conversion to non-forest (*3*, *5*). However, our results suggest a trade-off: reducing wildfire impact by preserving the stand works against the compositional turnover needed to re-establish a fully fire-adapted forest.

This interpretation is supported by our result that treatments increase tree survival most for species with low fire-tolerance. Because those fire-intolerant species would otherwise experience disproportionately high mortality during surface fires, treated stands can retain a larger fraction of their fire-maladapted pre-fire overstory. We interpret this as the mechanism behind the missing shift towards fire-tolerant species that a self-sustaining, fire-adapted forest would require: by preserving the stand, treatments also preserve its composition. The underlying problem, we argue, is that treatments restore a disturbance to an altered community, rather than restoring a community that is naturally in equilibrium with its own disturbance regime. Historically, many western U.S. forests experienced repeated low to medium intensity fires that continually shaped stand composition over decades by selecting for fire-tolerant species and leading to structural heterogeneity and low density stands (*8*, *9*). Treatments, by contrast, reduce fuel and stem density only temporarily, and appear neither to shift overstory composition enough nor to create the gaps that seedlings of fire-tolerant species require for germination (*29*). By making the altered forest more resistant, treatments protect precisely the composition that restoration aims to change.

This argument is further supported by the crown-fire observations in untreated stands. Untreated stands that subsequently burned as crown fires show substantially stronger shifts towards fire-tolerant regeneration, possibly because surviving fire-tolerant seed sources coincided with the high-light conditions created by canopy removal. If this interpretation is correct, treatments that sufficiently open the canopy—for example through intensive thinning combined with prescribed burning—may achieve similar regeneration without stand-replacing fire. Another possibility is that the observed composition reflects a mixture of surviving advanced regeneration and new post-fire recruits. If surface fire retains more of the pre-fire seedling cohort than crown fire, its regeneration layer may more strongly preserve the shade-tolerant, fire-intolerant composition associated with fire-excluded stands.

Treatment effectiveness varies substantially across climatic conditions, but the patterns differ between the resistance and regeneration component. Within forest types, tree survival benefits increase towards drier plots, suggesting that treatments are particularly effective where drought stress amplifies wildfire impacts. In contrast, shifts in seedling composition towards fire-tolerant species, although never distinguishable from zero, tend to be stronger on wetter plots. These contrasting patterns suggest that maximizing short-term resistance and promoting forest restoration may require different spatial strategies. Locations where treatments most effectively reduce wildfire impact are not necessarily those where they most strongly promote fire-adapted regeneration. Optimizing restoration may therefore require objectives and treatment designs that differ among regions rather than relying on a single management strategy.

Several limitations of the framework should be kept in mind. First, FIA data do not distinguish mechanical treatments (e.g. thinning) from prescribed burns, which work differently and have been shown to be most effective in increasing tree survival when used together and it also does not record treatment intensity (*12*). Second, FIA treatment records encompass all forms of tree removal, including commercial harvests, so we cannot identify harvests that removed large, fire-tolerant trees. Such harvests would remove fire-tolerant seed sources, and thus would impede fire-adapted regeneration. This would bias our estimates towards no effect rather than inflate them. Third, because our seedling analysis is restricted to plots with regeneration, it characterizes seedling composition rather than recruitment density. Treatments may increase fire-tolerant seedling abundance without strongly altering relative composition. Finally, because fires with minimal impacts are not recorded by FIA, our analysis excludes the least damaging fire events. Future work should further investigate how treatment intensity, spatial arrangement and different combinations of treatments affect treatment outcomes at scale.

## Conclusion

Our findings suggest a tension between maximizing short-term resistance and promoting long-term resilience in western U.S. forests. By protecting existing stands that are altered by a century of fire exclusion and management for timber production, treatments also preserve overstory communities that remain dominated by fire-intolerant species, limiting the compositional turnover through which recurrent fire historically maintained fire-adapted forests.

Achieving both objectives of treatments - fire hazard reduction and restoration of fire-adapted forests - may therefore require treatment strategies designed not only to reduce fuels but also to promote compositional change. This could include approaches that selectively reduce fire-intolerant overstory, create gaps in forests that allow for germination of shade-intolerant fire-adapted species, or combine repeated treatments with prescribed fire. We however posit that any spatial targeting faces a further complication: treatments do the most for tree survival on the driest sites, but that is not where they do the most for fire-tolerant regeneration.

As wildfires become increasingly severe under climate change, fuel treatments will remain a critical tool in reducing fire hazard in the short-term. The ultimate goal, however, is not only forests that survive a single fire, but forests capable of sustaining themselves and being resilient under recurrent fire without repeated treatments. To achieve this goal, our study suggests that current treatment protocols need to change.

## Acknowledgements

TM acknowledges funding from the La Kretz Center for California Conservation Science. CS acknowledges funding from the ORCA Acceleration Grant Wildfire Research Funding program from UCLA. The authors thank Malcolm North, Theresa Feo and Alex Hall for comments on a previous version of this manuscript.

## Data and Code availability

Only publicly available data was used in this study. FIA data was downloaded via the FIA datamart (https://research.fs.usda.gov/products/dataandtools/fia-datamart). Climate data were taken from WorldClim version 2.1 (https://worldclim.org/data/worldclim21.html) and TerraClimate (http://thredds.northwestknowledge.net:8080/thredds/fileServer/TERRACLIMATE_ALL/climatology/TerraClimate_19912020_def.nc). Code will be made available in a public repository after acceptance of the manuscript.

## Materials and Methods

### Study region and data

We used data from the U.S. Forest Service Forest Inventory and Analysis (FIA) database (*31*) (last access: February 5th, 2026) for 11 states in the western United States (Arizona, California, Colorado, Idaho, Montana, Nevada, New Mexico, Oregon, Utah, Washington, Wyoming). These states include the majority of dry fire-prone forest area in the western United States. Plots span forest types whose historical fire regimes range from frequent low- and mixed-severity (ponderosa pine, Sierra mixed conifer, Douglas-fir, oak-hardwood, pinyon-juniper) to infrequent stand-replacing fires (lodgepole pine, fir, aspen). To accommodate for differences among forest types, forest type enters every model as a random intercept. Forest-type-specific treatment contrasts are further estimated in separate models (see below).

### Data extraction and filtering

We extracted plot information from the PLOT table and information about treatments and fire history from the COND table. Tree and seedling composition were extracted from the TREE and SEEDLING tables, respectively. Only sampled plots containing at least one accessible forest land condition on the plot (PLOT_STATUS_CD = 1 and COND_STATUS_CD = 1) were considered. As plots could contain several conditions, plot-level summaries were calculated by weighting slope, southwestness and site productivity class with condition area and taking forest type and time since fire from the plot’s largest condition and classifying a plot as treated or as crown fire if any of its conditions was classified as such. These plot-level summaries were used for the propensity score, matching and inverse probability weighting, the fire-severity model and all seedling composition models in which seedling counts were summed over all conditions so that one observation is one plot. In the tree survival models, each tree carried the covariates, treatment and fire classification of the condition it was recorded in.

Only inventories measured in 2000 or later were included in the analysis, as this is the year when the FIA database was standardized and inventories hence comparable with respect to sampling procedure and timing of re-inventories. Only plots in which a fire occurred were selected based on the primary disturbance code (DSTRBCD1 = 30, 31 or 32). This was done as FIA records the disturbance judged most important for a condition as the primary disturbance code, so a fire appearing only in the second or third code was not the dominant disturbance at that plot, and this study focused solely on fire effects. A disturbance is noted in the FIA database when the area affected by it exceeds 1 acre in size and mortality or damage of at least 25% of trees in the condition is found. Code 31 records surface (ground) fire and code 32 crown fire. Code 30 records fire without specifying its type. Because code 30 was used only outside CA, OR and WA, the three states where most crown fires occur, we classified these plots as surface fire. The time since fire was calculated as the time between the fire record and the inventory year (MEASYEAR − DSTRBYR1).

Forest types were grouped into broader categories from the FIA forest-type codes (FORTYPCD), see Supplementary Table S2. Plots whose forest-type code fell outside any of these groups, and plots with a missing code, were excluded, as were hemlock–cedar plots, as they were not the focus of this study. Nonstocked stands (FORTYPCD 999) were retained. Forest type was taken from the post-fire inventory. Plots that were classified as nonstocked after a fire but were assigned a forest type other than “nonstocked” before the fire were assigned the forest type given before the fire, as it was assumed these plots fully burned and could hence not be assigned a forest type after the fire. Plots that were classified as nonstocked after a fire and had no forest type assigned before the fire either but had a FLDTYPCD indicating a forest type that was given by the field crew doing the inventory were assigned that FLDTYPCD. Plots missing key site variables (slope, elevation, site productivity class) or coordinates were excluded, as were plots for which no WorldClim or TerraClimate climate values (see below) could be extracted. In both analyses, plots were finally restricted to the region of propensity-score overlap (see section “Propensity score and positivity”), which removed plots with estimated propensity below 0.01 or above the 99th percentile of the propensity distribution.

### Treatment definition

Burned plots were considered treated when a treatment record was noted in the primary treatment field (TRTCD1) to occur in or before the year of the fire in the same plot. The analysis record for each plot is always the first post-fire inventory. Treatment information was carried forward to it from earlier inventories. FIA plots are measured on a ten-year cycle, so treatments applied before a fire are sometimes recorded in an earlier inventory than the fire itself. We therefore traced each plot back through its previous-plot records across up to two earlier inventory cycles and used the most recent treatment recorded in the primary treatment field (TRTCD1) of those records. A treatment was only counted when it occurred within 15 years before the fire and for treated plots we recorded the number of years between treatment and fire for use in the temporal analysis (see section “Tree survival model”). Only plots with treatment codes 10 (i.e. cutting, defined as “*the removal of one or more trees from a stand*” in the FIA database manual) and 20 (i.e. site preparation, defined as “*clearing, slash burning, chopping, disking, bedding, or other practices clearly intended to prepare a site for either natural or artificial regeneration”* in the FIA database manual) were used as treated plots. Plots with post-fire planting (treatment code 30) in the current or two previous inventories were excluded because we wanted to investigate only the effect of the natural ecosystem response to wildfire disturbance. The FIA database categorizes a plot as treated if the treated area exceeds 1 acre in size.

### Tree dataset filtering

Trees were retained when measured between 1 and 10 years after the fire, because this is the time interval in which plots are measured. Only live trees (STATUSCD = 1) and trees recorded as dead with fire as the mortality agent (STATUSCD = 2, AGENTCD = 30) were analysed. Trees that died from other agents were excluded so that the outcome only considers fire-caused mortality. Trees removed by cutting (STATUSCD = 3) were excluded. Trees that entered the inventory as regrowth since the previous measurement (RECONCILECD 1 or 2) were excluded, as they were not standing at the time of the fire, as were trees recorded as fire-killed but whose mortality year preceded the fire year. Trees smaller than 1 inch diameter were excluded, with diameter taken from the current measurement (DIA) or, where missing, from the previous measurement (PREVDIA). Species were retained when represented by at least 100 trees, of which at least 25 occurred on treated plots. The final number of plots and trees for treated and untreated conditions, as well as surface and crown fire categories can be found in Supplementary Table S3.

### Seedling dataset filtering

Seedling records were retained when the plot was measured between 2 and 10 years after the fire. A lower threshold of two years was chosen so that post-fire establishment is detectable, and a maximum threshold of ten years was chosen so that the observed cohort still reflects initial post-fire regeneration. Seedling counts (TREECOUNT) were aggregated to the plot level within three fire-tolerance classes (low, moderate, high), so that one observation is one plot. Species were retained when present on at least 10 plots with at least 30 seedlings in total, and only species for which a fire-tolerance classification could be assigned were used. Plots on which no seedlings of any retained species were recorded were excluded, so the compositional analysis describes which species regenerated on plots where regeneration occurred, and not whether regeneration occurred at all. This excluded 57 plots. The final number of plots and seedlings for treated and untreated conditions, as well as surface and crown fire categories can be found in Supplementary Table S4.

### Species fire tolerance classification

Species were classified into low, moderate or high fire-tolerance based on species descriptions in the Fire Effects Information System (FEIS) (*32*). All classifications can be found in Supplementary Table S5.

### Climate data

Mean annual temperature (MAT) and mean annual precipitation (MAP) were obtained for each plot from the WorldClim version 2.1 bioclimatic dataset (*33*) at a resolution of 2.5 arc-minutes (approximately 4.5 km) and climatic water deficit (CWD) was obtained from TerraClimate (*34*) at a comparable resolution (1/24 degrees corresponding to about 4 km). CWD was calculated as the annual sum (mm/yr). This resolution was assumed to be the right scale matching the coordinate fuzziness of the FIA database (approximately 1 mile/1.6 km). MAT, MAP and CWD were used in the propensity-score and matching models to adjust for climatic differences between treated and untreated plots. They were additionally included as covariates after standardization (z-score) in the tree survival and seedling composition shift models and the interaction of CWD with treatment was used in an additional model to test whether treatment efficacy depends on climate.

### Causal inference framework

Fuel treatments are not assigned randomly but are more likely to be deposited in specific forest locations that are for example more accessible or more likely to be subject to high severity fires due to climate, topography or forest structure (*35*). Hence, any estimate of a causal effect of treatments on tree survival and seedling composition shift needs to address the influence of these confounders. Furthermore, treatments may act through a direct pathway in increasing tree survival or shifting seedling composition for example through a reduction in competition for resources among trees and seedlings, but they may also act over an indirect pathway by reducing fire-severity which then increases tree survival or shifts seedling composition. Those pathways need to be disentangled to identify treatment benefits on tree survival and seedling composition shift at the surface fire conditions they are intended to promote.

We start from a directed acyclical graph (see Fig. 1c) that encodes the proposed causal pathway describing how treatments increase survival and shift seedling composition (*36*, *37*). Next to the direct pathway and the indirect pathway over lowering fire severity we have three blocks of confounders that represent pre-treatment plot-level variables all of which influence the likelihood that treatments are placed in a plot but that also themselves influence crown fire probability and the outcome (tree survival, seedling composition shift) (*35*, *38*). These three blocks are topography (represented by elevation, slope and southwestness), climate (MAT, MAP, CWD) and stand structure/site capability (forest-type group, site productivity class).

For correct causal identification of treatment effects, the following assumptions need to be met, and for each one we state how we addressed them:

1. No unmeasured confounding: Treated and untreated plots need to be comparable given the three confounder blocks. We assessed sensitivity to unmeasured confounding with E-values and negative-control outcomes (*39*, *40*).
2. Positivity: Every plot must have a non-zero probability of being treated or untreated. We enforce this by calculating a propensity score, trimming the region without common support and removing any forest type or state observed only in either the treated or untreated plots (*41*).
3. Consistency: The categories “treated” and “untreated” must be consistently defined (*42*). This is achieved by using the categorization of the FIA database as described above (see “Treatment definition”).
4. No interference: Treatments are assigned at plot level and a treatment in one plot should not affect causal effects (i.e. tree survival and seedling composition shift) in another plot (*43*). As FIA plots are located roughly 5 km apart from each other (*44*), treatment of one plot is unlikely to change fire behaviour or regeneration in another. We modelled the remaining dependence with plot-level random effects and a spatial field in the Bayesian models.

### Statistical analyses

To address whether treatments increase short-term resistance of forests to wildfire, we modelled the effect of treatments on tree survival and crown fire probability. To address whether treatments also lead to fire-adapted regeneration of forests we modelled the effect of treatments in shifting the species composition of seedlings towards seedlings of fire-tolerant species. To estimate treatment effects on tree survival and seedling composition shift we used a hierarchical Bayesian framework in R-INLA v25.10.19. Writing of code was assisted by Claude Opus v4.8, v5 and v5.5. All code was checked and verified by the authors. For both models we report two causal quantities, which are the total effect of treatments on tree survival and seedling composition shift and the controlled direct effect. The total effect is the effect of treatments along every pathway, including the one over reduced fire severity. The controlled direct effect is the effect of treatments when holding fire-severity fixed, i.e. it explains the treatment effect over mechanisms other than reducing fire-severity (see Fig. 1c).

### Propensity score and positivity

To ensure positivity (see point 2 in section “Causal inference framework”), a plot-level logistic propensity score was estimated from the pre-treatment covariates elevation, slope, southwestness, site productivity class, the climate variables, forest type and state. The model is fitted at the plot level, because treatments are applied to whole plots. First, any forest types or states that either had only treated or untreated plots were removed to ensure every forest type and state has both treated and untreated plots. Second, the propensity score was fitted and plots with a score below 0.01 or above the 99th percentile of all fitted scores were trimmed. Third, the check for forest types and states that only had treated or untreated plots was applied again because trimming can leave a forest type or state with either. Fourth, the propensity score was refitted on this final sample and the fitted scores were bounded to the range [0.01, 0.99]. This refitted score was used for weighting, matching and AIPW (see section “Robustness”). After this procedure, 386 of 4,015 plots in the tree survival analysis and 281 of 2,042 plots for the seedling composition shift analysis were excluded. The retained region is the common support on which every estimate rests (see Supplementary Figures S7a and S8a for tree and seedling data, respectively).

### Tree survival model

Survival was modelled as a binary response following a binomial distribution with a logit link. Our primary model (Model A) reports the total effect of treatments on tree survival. The linear predictor included the treatment effect as well as an explicit intercept term, a quadratic effect of diameter at breast height (DBH), time since fire and covariates for the site (slope, southwestness, elevation, site productivity class, MAT, MAP and CWD). A quadratic effect for DBH was included because tree size was assumed to non-linearly influence tree survival, as increasingly larger trees are not expected to be more resistant to fire. Southwestness was calculated from the aspect of the plot as:

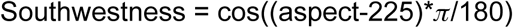

Plots with an aspect value of 0 or a slope of 0 were treated as missing for this calculation and subsequently the southwestness was set to 0 for those plots. The tree diameter was taken directly from the current measurement when available (column DIA) but filled with the diameter from a previous measurement (column PREVDIA) when unavailable. All continuous covariates were standardized (z-scored) before analysis. Plot-level covariates (elevation, slope, site productivity class, southwestness, time since fire, MAT, MAP, CWD) were standardized with the mean and standard deviation across plots, tree diameter was standardized across trees. The hierarchical structure is considered through Gaussian random intercepts for the plot, state, forest type and species. Additionally, species-specific random slopes for treatment were used because different species were expected to differently respond to treatments, especially with respect to different fire-tolerance of species. Species-specific random slopes for DBH were included because the size-mortality relationship was also assumed to differ depending on species, as large trees of fire-tolerant species are expected to withstand fire more successfully than large trees of fire-intolerant species. To account for residual spatial autocorrelation, a spatial Gaussian random field over sample locations was included using a stochastic partial differential equation (SPDE) field on a mesh with a maximum inner edge of 50 km, a maximum outer edge of 200 km, a 10 km cutoff and inner and outer extensions of 50 and 100 km. We used penalized complexity priors for the random-effect variances and for the range and standard deviation of the spatial field, which shrink each term towards zero unless the data support a stronger effect. Fixed effects received weakly informative Gaussian priors, N(0, 2.5²) for the main effects and N(0, 1.5²) for the quadratic DBH term and for all interactions, with a N(0, 32²) prior on the intercept.

We then fitted another model (Model B) that adds crown fire and its interaction with treatment and hence estimates the controlled direct effect at surface severity (the treatment coefficient) and under crown severity (treatment plus interaction coefficient).

We further investigated effects of climate, treatment longevity and forest type on treatment benefits with a series of additional models. These models are extensions of Model B, so the treatment effects they report are controlled direct effects at surface fire. First, we wanted to test whether aridity changes the treatment effect. To separate the effect of aridity from differences in forest types, we performed a Mundlak decomposition. For this, we split CWD into each forest type’s mean (between-forest-type component) and each plot’s deviation from that mean (within-forest-type component), both scaled by the same standard deviation across plots of the analysed forest types and both interacted with treatments. Only the within-forest-type component, which compares drier with wetter plots within a forest type, is interpreted because the between-forest-type component cannot be separated from other differences among forest types and only serves as an adjustment. Each forest type had its own random deviation in the treatment effect. We additionally investigated treatment longevity in a separate model by allowing the treatment effect to vary with the time between treatment and fire using a second-order random-walk smoother over the number of years between treatment and fire (0-15 years, with one node per year). Because this term is only defined for treated trees, it is identified from the treated subset. Forest-type-specific decay was estimated by letting the treatment effect change linearly with treatment age, with independent forest-type random effects on both the treatment effect and its slope. The Mundlak decomposition and the forest-type-specific decay model were fitted only to forest types with at least 10 treated plots, because per-type estimates resting on fewer treated plots are unreliable. Pooled models retained all forest types.

Fire severity was modelled at the plot level on burned plots only, with crown fire (FIA DSTRBCD1 = 32) versus surface fire (FIA DSTRBCD1 = 30, 31) as a binary response using a binomial likelihood with a logit link. The linear predictor included an explicit intercept, a fixed effect for treatment, and the same confounder set used for the propensity score model which were topography and site (elevation, slope, southwestness, site productivity class), climate (MAT, MAP, CWD) as well as forest type and state, so that all estimates rest on a common adjustment set. Continuous confounders were standardized (z-scored) within this subsample. Residual spatial dependence was modelled with a Gaussian random field over plot locations on the same SPDE mesh used for the survival models. Species terms were not included, as severity is a plot-level property. Fixed effects and the intercept received N(0, 2.5²) priors.

INLA returns marginal posterior distributions for all fixed and random effects. For the tree survival and fire-severity models, fixed effects are estimated on the log-odds scale, because they are binomial models with a logit link function. These were converted to odds-ratios (OR) that are easier interpretable as they present the change in survival or crown fire odds due to a fixed effect. 95% credible intervals (CrIs) were calculated as the 2.5th and 97.5^th^ percentiles of the exponentiated posterior marginal distributions. Odds ratios are reported as the exponentiated posterior mean of the log-odds coefficient. Forest-type-level treatment effects were computed from 2,000 draws of the joint posterior, so that intervals carry the covariance between the fixed coefficient and the forest-type deviation. Species-level effects were approximated from Model B as the fixed coefficient plus the species deviation with variances added (±1.96 SD). Fire-tolerance-class odds ratios in Fig. 2b were obtained by inverse-variance pooling of species-level effects. An OR > 1 indicates an increased tree survival or probability of a crown fire event, whereas an OR < 1 indicates a decrease in probability for the event. As all continuous covariates were standardized (z-scored), the OR represents the multiplicative change in odds associated with a one standard deviation increase in the covariate.

### Seedling composition shift model

Counts were aggregated to plot level by tolerance class, so each plot contributed three observations. Seedling counts were modelled with a negative binomial likelihood using the plot’s total seedling count as an offset, following a multinomial–Poisson formulation. The linear predictor is therefore on the log-proportion scale, and predicted composition follows from a softmax transformation across the three classes:

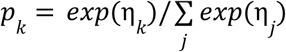

where η*_k_* is the linear predictor for tolerance class *k*.

Like in the tree survival model one primary model (Model A) reports the total effect of treatments on seedling composition shift. The predictor included one intercept per tolerance class, a class-specific treatment effect, time since fire and covariates for the site (slope, southwestness, elevation, site productivity class, MAT, MAP and CWD) each interacted with tolerance class. Southwestness was calculated as in the tree survival model. All continuous covariates were standardized (z-scored) before analysis. Gaussian random intercepts were included for plot, state and forest type. Forest type and state additionally entered as tolerance-class-specific random deviations (for the moderate and high classes relative to low). Residual spatial structure in seedling abundance was modelled with an SPDE field on a mesh with a maximum inner edge of 50 km, a maximum outer edge of 200 km, a 10 km cutoff and inner and outer extensions of 50 and 100 km. All variance components received penalised-complexity priors, the same as in the survival model. Main effects, including the class-specific treatment effects, received N(0, 2.5²) priors and all treatment interactions N(0, 1.5²) priors, as in the survival model, while the class intercepts received N(0, 10²) priors.

We then also fitted another model (Model B) that adds crown fire and its interaction with treatment and hence estimates the controlled direct effect at surface severity (the treatment coefficient) and under crown severity (treatment plus interaction coefficient).

Because predicted composition is a nonlinear function of several correlated coefficients, all predicted compositions were obtained from 2000 draws from the joint posterior. For the total and controlled direct effects, for each draw, the class-specific linear predictor was computed for every observed plot under each treatment × severity combination, using that plot’s own covariates and forest-type and state deviations. The softmax was applied per plot and compositions were averaged over plots (marginal standardisation). We report posterior medians and 95 % credible intervals. To test whether the compositional shift changes over time as seedlings germinate we extended Model B by adding time since fire interacted in a three-way interaction with tolerance class and treatment. Predictions were made at surface fire, mean values of the remaining covariates and for an average forest-type and state (random deviations set to 0). Time since fire was predicted over its observed range (2–10 years). Because time since fire varies between rather than within plots, the temporal pattern is a cross-sectional gradient in post-fire stand age rather than an observed regeneration trajectory.

To test an effect of aridity on treatment effects in shifting seedling composition we did, as in the tree survival model, a Mundlak decomposition to separate aridity effects from differences across forest types. To do so, CWD was split into a between-forest-type component (each type’s mean relative to the global mean) and a within-forest-type component (each plot relative to its own type mean), both scaled by the same global standard deviation so that the coefficients are directly comparable. Extending Model B, both entered as tolerance-specific covariates and in three-way interactions with treatment and tolerance class. Analyses in this model were restricted to forest types with at least 5 treated plots to ensure robustness. Because these interactions are estimated from limited within-type CWD spread, we placed weakly informative N(0, 1.5²) shrinkage priors on all six. The decomposition coefficients are reported directly from the posterior marginals. The compositional shift along the within-forest-type CWD component was computed from posterior draws for the same settings as for time since fire.

### Robustness

We used several approaches to test the robustness of our results with respect to uneven balance of selection of treated plots as well as the used methodology and unmeasured confounding.

All these methods start with the propensity score as described in the section “Propensity score and positivity” above. The tree-level survival analysis and the seedling composition analysis each use their own score, fitted to the same site and climate covariates. The balance was summarised as the absolute standardized mean differences against a threshold of 0.1 using the raw pooled standard deviation as the common fixed denominator per covariate. Treated plots are generally lower in elevation, more productive and drier than untreated plots (see Supplementary Figures S7b and S8b).

With the propensity score we re-estimated the total effect in three ways. First, with inverse probability weighting (IPW) where each plot gets a stabilized weight from the inverse of its propensity score so that the treated and untreated groups become comparable. Second, with propensity score matching, pairing each treated plot with the untreated plot with the closest propensity score (1:1 nearest neighbour without replacement, caliper of 0.2 standard deviations of the propensity score). For both, Model A was refitted with an otherwise unchanged specification, once on the matched sample and once on the full sample with the IPW weights entering the likelihood, so that only the sample or its weighting differs from the main estimate. Third, we used augmented inverse probability weighting (AIPW), a doubly robust approach that combines the propensity score with separate outcome models for treated and untreated plots and which gives a valid estimate as long as at least one of the two models is correct. For tree survival, the outcome models were logistic regressions of tree survival. For seedling composition, they were binomial regressions of each tolerance class’s share of the plot’s seedlings, and the three class estimates were rescaled to sum to one. Neither included the random effects or the spatial field. In both analyses the outcome models used the full propensity-score covariate set plus time since fire. Because treatment is a plot-level variable but survival is measured on individual trees, the uncertainty of the tree survival AIPW estimate was calculated with standard errors clustered at the plot level. For the seedling composition AIPW estimate, uncertainty came from 1,000 cluster-bootstrap replicates over plots, with the propensity score refitted in each replicate.

### Sensitivity

We asked how strong an unmeasured confounder would need to be to overturn our result. For this we calculated the E-value for the total effect of treatments on tree survival (Model A for tree survival model) which can be interpreted as the minimum strength of association, on the risk-ratio scale, that an unmeasured variable would need to have with both treatment assignment and tree survival, conditional on the measured covariates, to explain the effect away (*39*) and compared it with the strength of the confounders we had actually measured. Because survival is not a rare outcome, odds ratios were converted to approximate risk ratios via the square-root transformation (*45*) before computing E-values. The E-value was 2.72 for the point estimate and 1.67 for the lower credible limit (Table S1). We complemented this sensitivity analysis with a falsification test that would be expected to reveal residual confounding if there was still a systematic difference between treated and untreated plots beyond what our models account for. We tested for a treatment effect on time since fire (the interval between the fire year and the FIA revisit) as this cannot be caused by a treatment applied before the fire. The test regressed time since fire on treatment using weighted least squares with the same stabilized, trimmed inverse-probability weights used above, adjusting for the remaining topographic and site variables (elevation, slope, southwestness, site productivity class, MAT, MAP, CWD) and for forest-type and state. The regression was fitted at the plot level rather than on individual trees, because the tested variables are plot attributes. After weighting, treatment did not predict time since fire in the survival dataset (β = 0.050 SD, p = 0.48) or in the seedling dataset (β = 0.088 SD, p = 0.39), indicating no detectable systematic difference between treated and untreated plots when looking at time since fire.

## Supplementary Material

**Supplementary Table S1.**
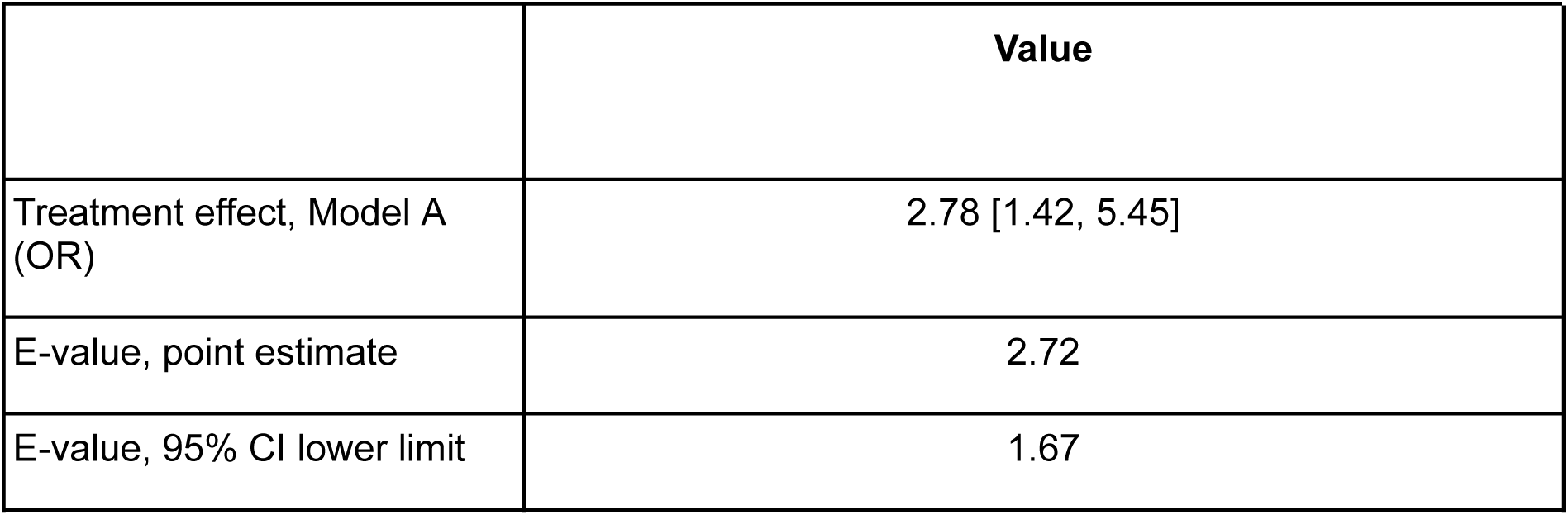
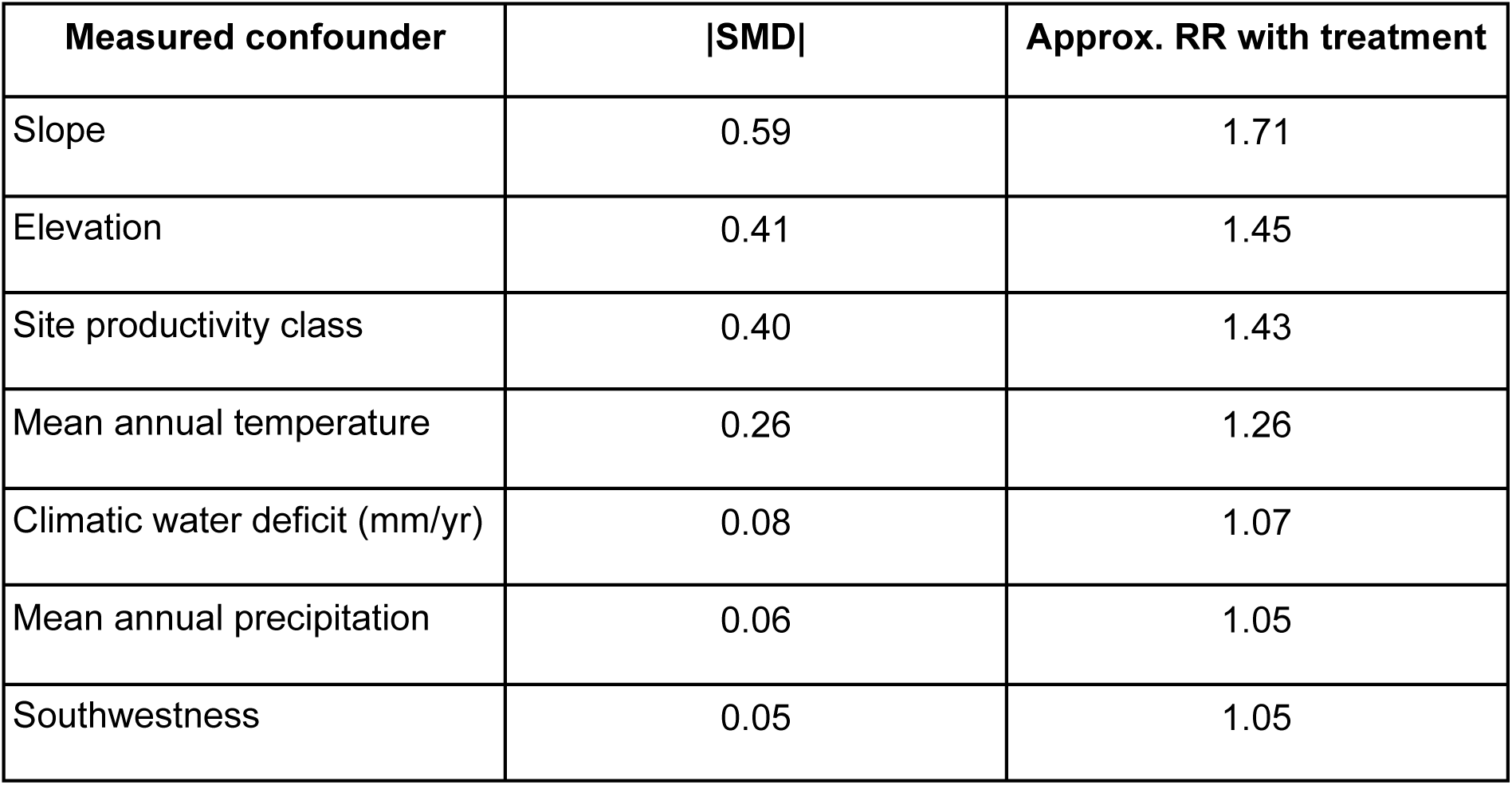
E-values for the total treatment effect on tree survival (Model A). Because survival is not a rare outcome, odds ratios were converted to approximate risk ratios by the square-root transformation. Standardised mean differences (SMD) were placed on the same scale as exp(π/(2√3) × |SMD|). SMDs were calculated on the data before trimming.

|  | **Value** |  |
|---|---|---|
| Treatment effect, Model A (OR) | 2.78 [1.42, 5.45] |  |
| E-value, point estimate | 2.72 |  |
| E-value, 95% CI lower limit | 1.67 |  |
| **Measured confounder** | **\|SMD\|** | **Approx. RR with treatment** |
| Slope | 0.59 | 1.71 |
| Elevation | 0.41 | 1.45 |
| Site productivity class | 0.40 | 1.43 |
| Mean annual temperature | 0.26 | 1.26 |
| Climatic water deficit (mm/yr) | 0.08 | 1.07 |
| Mean annual precipitation | 0.06 | 1.05 |
| Southwestness | 0.05 | 1.05 |

**Supplementary Table S2.** Forest type code classifications. Taken from the FIA database user guide (Appendix D) version from 8/2025 (*46*).

| **Forest type code** | **Forest type group** |
|---|---|
| 182,184,185,369 | Pinyon-Juniper |
| 201-203 | Douglas fir |
| 221,222,224 | Ponderosa dry |
| 225-226 | Jeffrey pine |
| 241,362,365,366,367 | White pine-Limber |
| 260-271 | Fir |
| 280,281 | Lodgepole |
| 301,304,305 | Hemlock cedar |
| 321 | Western larch |
| 341,342 | Redwood |
| 371 | Sierra mixed conifer |
| 361,363,364,368 | Other conifer |
| 703,704,706,709,722,911,912 | Riparian hardwood |
| 901,902 | Aspen |
| 920-924,931,933-935,941-943,961,962 | Oak hardwood |
| 971-976 | Oak woodland |
| 999 | Nonstocked other |

**Supplementary Table S3.**
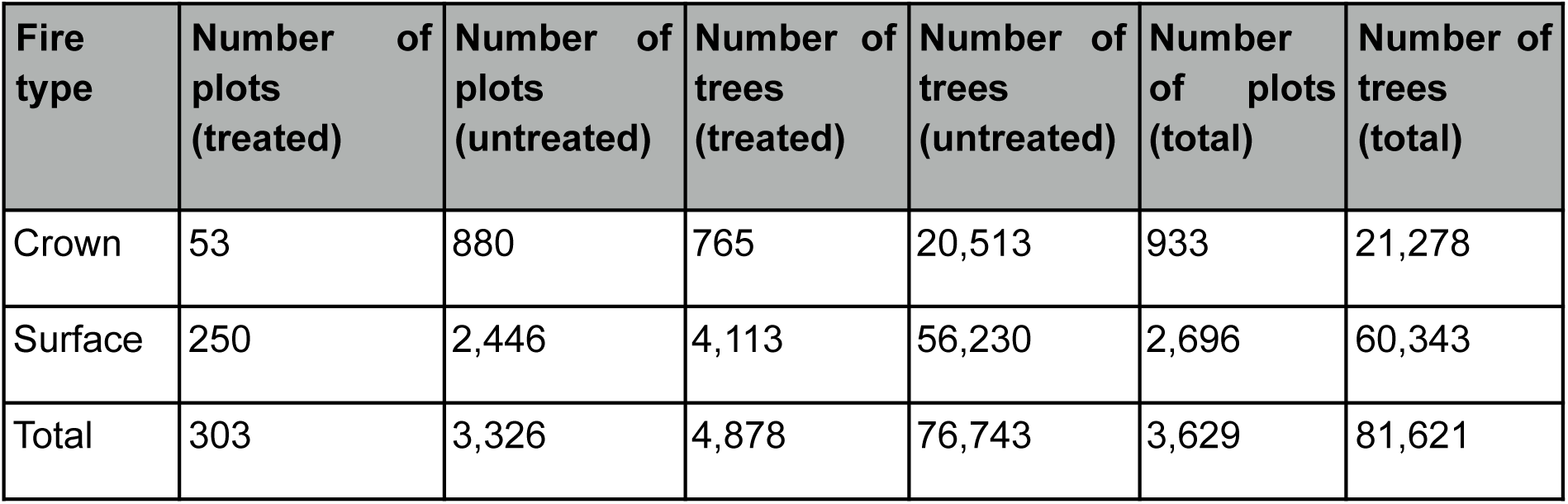
Number of plots and trees for various categories for the tree survival model.

| **Fire type** | **Number of plots (treated)** | **Number of plots (untreated)** | **Number of trees (treated)** | **Number of trees (untreated)** | **Number of plots (total)** | **Number of trees (total)** |
|---|---|---|---|---|---|---|
| Crown | 53 | 880 | 765 | 20,513 | 933 | 21,278 |
| Surface | 250 | 2,446 | 4,113 | 56,230 | 2,696 | 60,343 |
| Total | 303 | 3,326 | 4,878 | 76,743 | 3,629 | 81,621 |

**Supplementary Table S4.** Number of plots and seedlings for various categories for the seedling composition shift model.

| **Fire type** | **Number of plots (treated)** | **Number of plots (untreated)** | **Number of seedlings (treated)** | **Number of seedlings (untreated)** | **Number of plots (total)** | **Number of seedlings (total)** |
|---|---|---|---|---|---|---|
| Crown | 26 | 445 | 390 | 12,667 | 471 | 13,057 |
| Surface | 112 | 1,178 | 1,676 | 26,595 | 1,290 | 28,271 |
| Total | 138 | 1,623 | 2,066 | 39,262 | 1,761 | 41,328 |

**Supplementary Table S5.** Species were classified into low, moderate or high fire-tolerance based on species descriptions in the FEIS (*32*).

| **Common name** | **Scientific name** | **Fire tolerance** |
|---|---|---|
| Pacific silver fir | Abies amabilis | Low |
| White fir | Abies concolor | Low |
| Grand fir | Abies grandis | Low |
| Corkbark fir | Abies lasiocarpa arizonica | Low |
| Subalpine fir | Abies lasiocarpa | Low |
| Red fir | Abies magnifica | Low |
| Shasta red fir | Abies magnifica shastensis | Low |
| Noble fir | Abies procera | Low |
| Port Orford cedar | Chamaecyparis lawsoniana | Low |
| Alaska cedar | Callitropsis nootkatensis | Low |
| Arizona cypress | Hesperocyparis arizonica | Moderate |
| Baker cypress | Hesperocyparis bakeri | Moderate |
| Sargents cypress | Hesperocyparis sargentii | Moderate |
| Incense cedar | Calocedrus decurrens | Moderate |
| California juniper | Juniperus californica | Low |
| Alligator juniper | Juniperus deppeana | High |
| Western juniper | Juniperus occidentalis | Moderate |
| Utah juniper | Juniperus osteosperma | Moderate |
| Rocky Mtn juniper | Juniperus scopulorum | Moderate |
| One-seed juniper | Juniperus monosperma | Moderate |
| Subalpine larch | Larix lyallii | Moderate |
| Western larch | Larix occidentalis | High |
| Engelmann spruce | Picea engelmannii | Low |
| Blue spruce | Picea pungens | Low |
| Whitebark pine | Pinus albicaulis | Low |
| Bristlecone pine | Pinus aristata | Moderate |
| Knobcone pine | Pinus attenuata | Moderate |
| Pinyon (common) | Pinus edulis | Moderate |
| Lodgepole pine | Pinus contorta | Low |
| Coulter pine | Pinus coulteri | Moderate |
| Apache pinyon | Pinus engelmannii | Moderate |
| Limber pine | Pinus flexilis | Moderate |
| SW white pine | Pinus strobiformis | Moderate |
| Jeffrey pine | Pinus jeffreyi | High |
| Sugar pine | Pinus lambertiana | Moderate |
| Western white pine | Pinus monticola | Moderate |
| Ponderosa pine | Pinus ponderosa | High |
| Gray pine | Pinus sabiniana | Moderate |
| Single-leaf pinyon | Pinus monophylla | Moderate |
| Pinyon (border) | Pinus discolor | Moderate |
| Douglas-fir | Pseudotsuga menziesii | Moderate |
| Coast redwood | Sequoia sempervirens | High |
| Giant sequoia | Sequoiadendron giganteum | High |
| Pacific yew | Taxus brevifolia | Low |
| Western redcedar | Thuja plicata | Low |
| Western hemlock | Tsuga heterophylla | Low |
| Mountain hemlock | Tsuga mertensiana | Low |
| Bigleaf maple | Acer macrophyllum | Moderate |
| Red alder | Alnus rubra | Low |
| White alder | Alnus rhombifolia | Low |
| Arizona alder | Alnus oblongifolia | Low |
| Pacific madrone | Arbutus menziesii | Moderate |
| Giant chinquapin | Chrysolepis chrysophylla | Moderate |
| Tanoak | Notholithocarpus densiflorus | Low |
| Quaking aspen | Populus tremuloides | Low |
| Black cottonwood | Populus trichocarpa | Low |
| Narrowleaf cottonwood | Populus angustifolia | Low |
| California live oak | Quercus agrifolia | High |
| Arizona white oak | Quercus arizonica | High |
| Canyon live oak | Quercus chrysolepis | Moderate |
| Blue oak | Quercus douglasii | Moderate |
| Emory oak | Quercus emoryi | Moderate |
| Engelmann oak | Quercus engelmannii | Moderate |
| Gambel oak | Quercus gambelii | Moderate |
| Oregon white oak | Quercus garryana | High |
| California black oak | Quercus kelloggii | High |
| Interior live oak | Quercus wislizeni | Moderate |
| Silverleaf oak | Quercus hypoleucoides | Moderate |
| Gray oak | Quercus grisea | Moderate |
| Netleaf oak | Quercus rugosa | Moderate |
| California laurel | Umbellularia californica | Moderate |

## Supplementary Figures

**Supplementary Figure S1:**
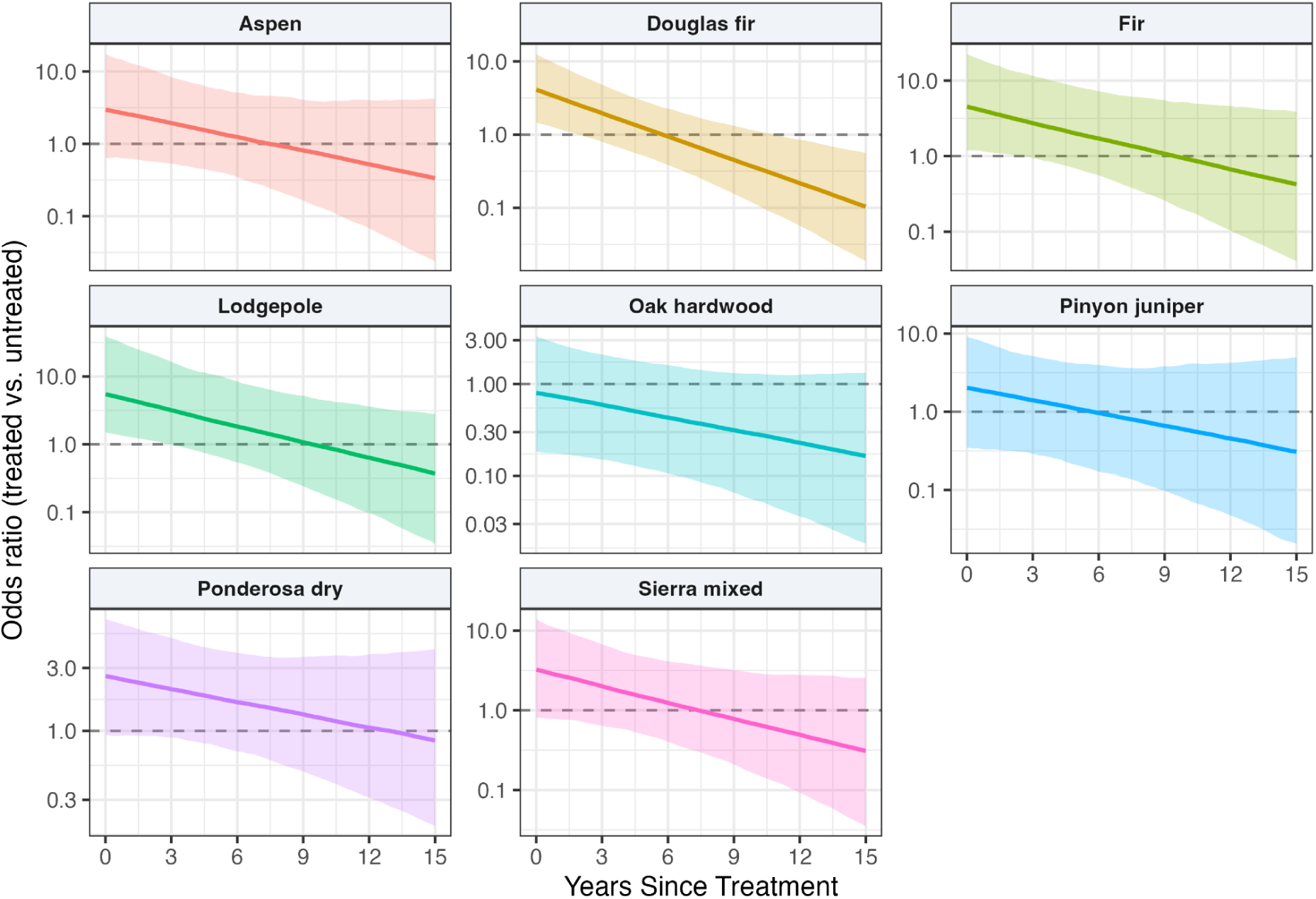
Treatment decay according to forest type. Dotted line indicates OR of 1, referring to no effect of treatments. Lines show posterior medians and bands 95% credible intervals, both based on 2,000 posterior draws.

**Supplementary Figure S2:**
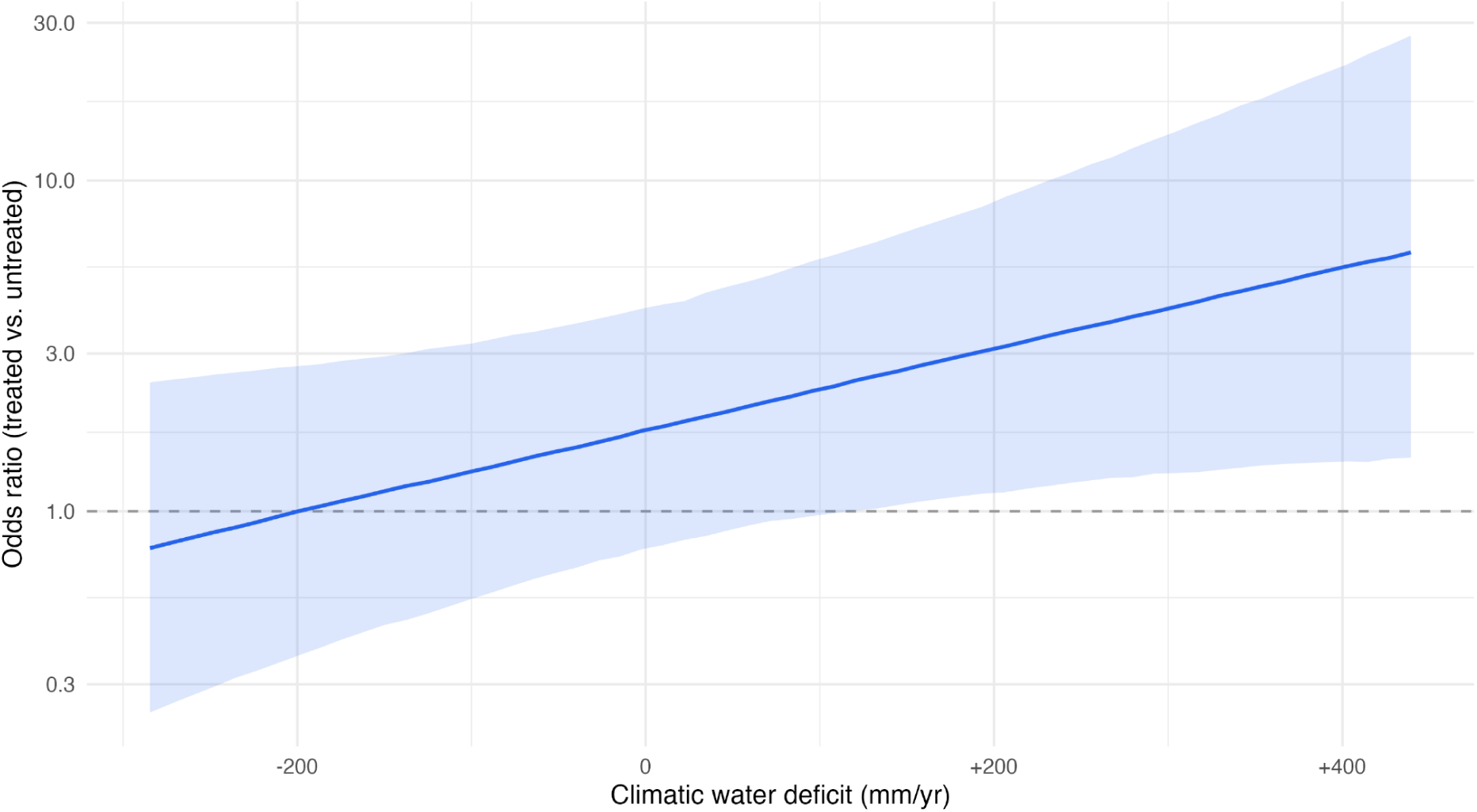
Within forest type CWD modulation of treatment effects on tree survival. The within forest type component shows a plot’s deviation from the forest type’s mean. The line shows the posterior median and the band represents the 95% credible interval, both based on 2,000 posterior draws. The within forest type component is based on the central 95% of plots.

**Supplementary Figure S3:**
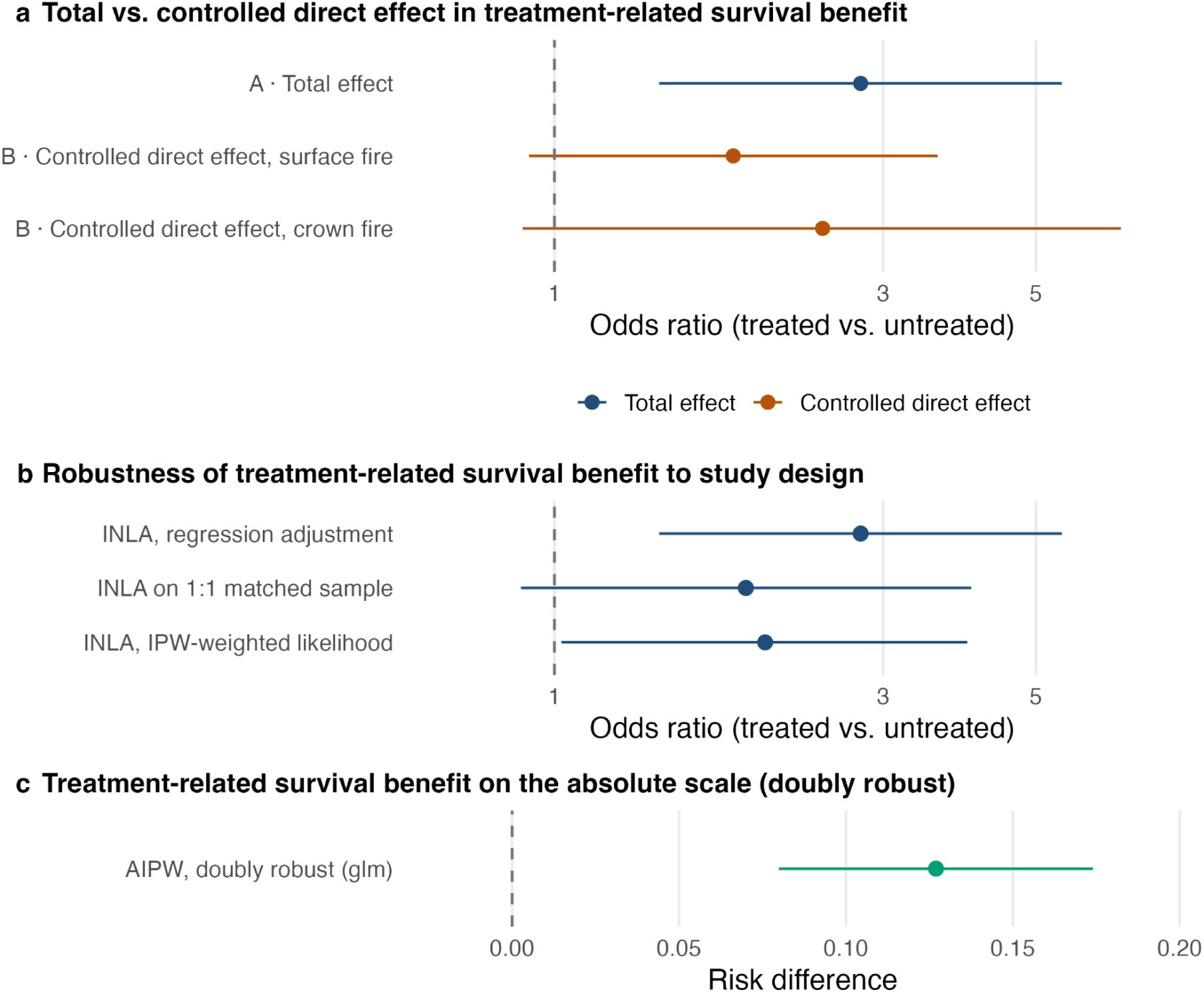
Robustness analysis for tree survival. (a) Total effect (Model A) and controlled direct effect at surface and crown fire (Model B) (b) Robustness of the total effect. Point estimates are medians and error bars in (a) and (b) represent 95% credible intervals, both based on 2,000 posterior draws. (c) Augmented inverse-probability-weighted (AIPW) estimate of the total effect, displayed as the marginal risk difference in survival probability. The bar represents 95% confidence interval. Dashed lines mark no effect. (a) and (b) show conditional ORs, (c) shows the population-averaged risk difference. Points in (a) as well as the regression adjustment, the IPW-weighted likelihood and the AIPW are based on the full trimmed sample consisting of 81,621 trees on 3,629 plots (303 treated, 3,326 untreated). The 1:1 matched sample is based on 12,158 trees on 604 plots (302 pairs of treated/untreated plots).

**Supplementary Figure S4:**
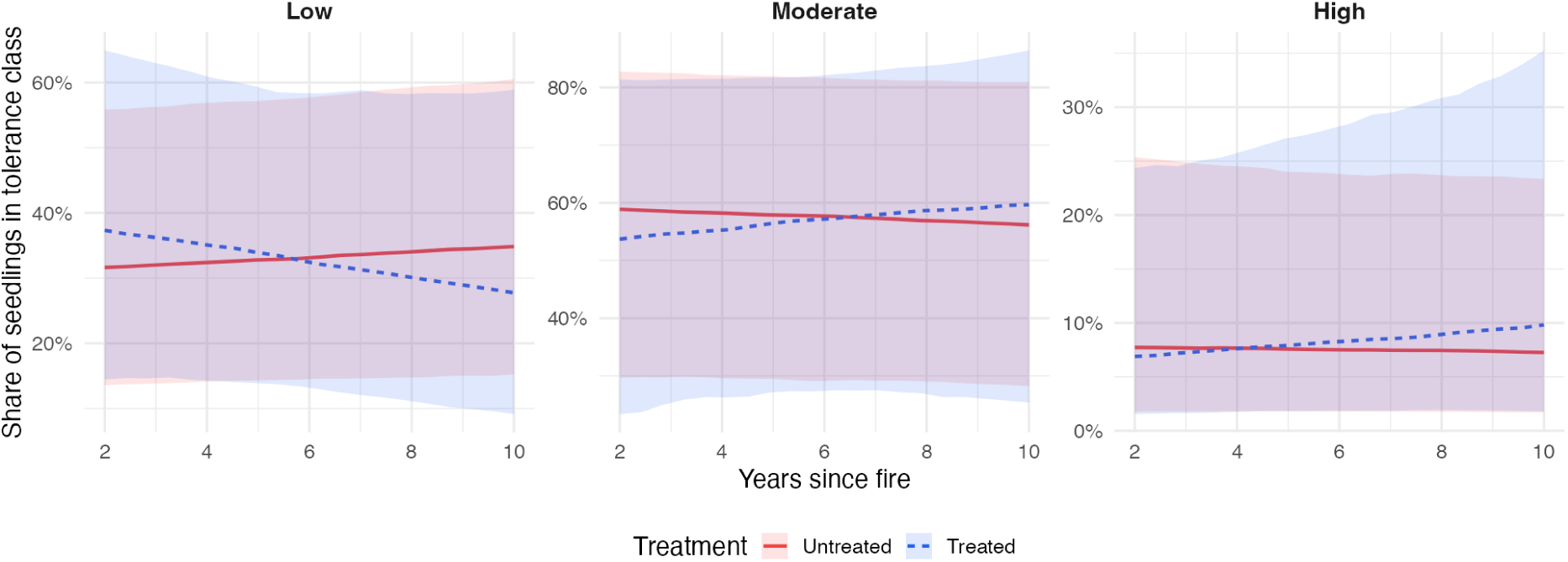
Predicted proportion of seedling composition per year after fire. Lines represent posterior medians and bands represent 95% credible intervals, both based on 2,000 posterior draws. Each plot is measured once, so the horizontal axis compares different plots at different times since fire rather than following plots through time. The window of 2-10 years is based on the FIA remeasurement interval of 10 years.

**Supplementary Figure S5:**
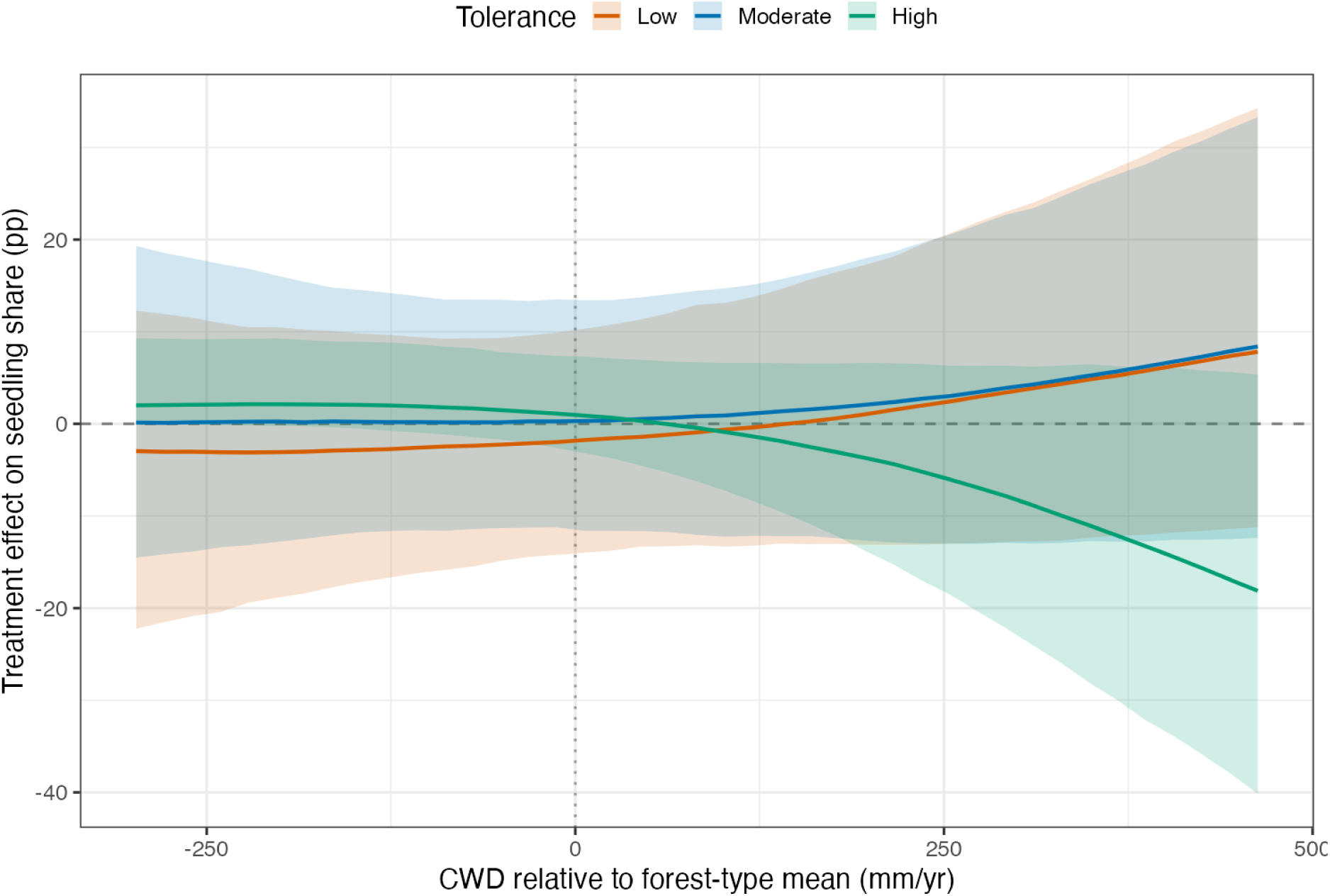
Within forest type CWD modulation of treatment effects on seedling composition shift. The within forest type component shows a plot’s deviation from the forest type’s mean. The within forest type component is based on the central 95% of plots.

**Supplementary Figure S6:**
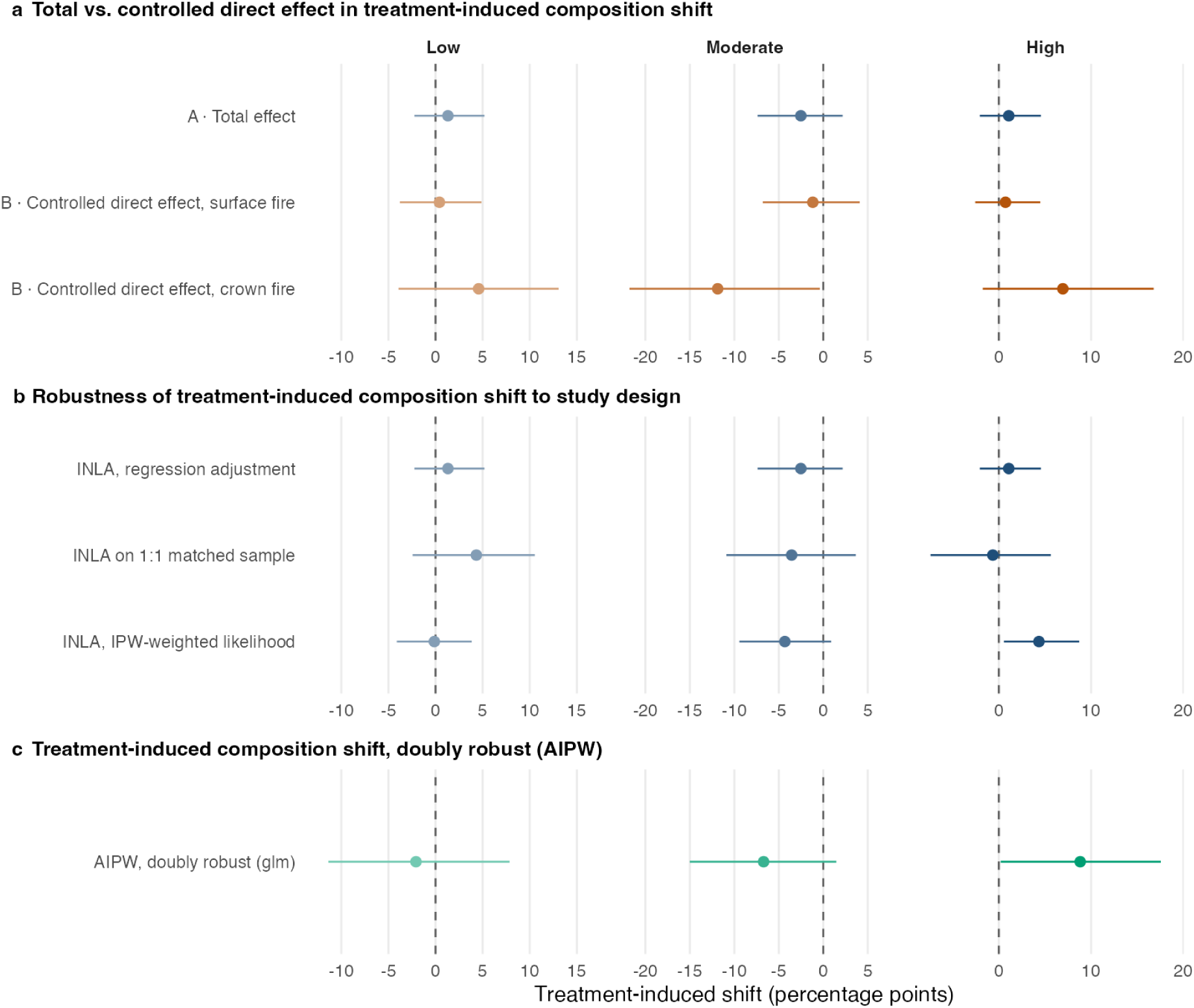
Robustness analysis for seedling composition shift. (a) Total effect (Model A) and controlled direct effect at surface and crown fire (Model B). (b) Robustness of the total effect. Point estimates are medians and error bars in (a) and (b) represent 95% credible intervals, both based on 2,000 posterior draws. (c) Augmented inverse-probability-weighted (AIPW) estimate of the total effect, displayed as the shift in seedling share. The bar represents 95% percentile intervals from a cluster bootstrap over plots (1,000 replicates). Dashed lines mark no effect. Points in (a) as well as the regression adjustment, the IPW-weighted likelihood and the AIPW are based on the full trimmed sample consisting of 41,328 seedlings on 1,761 plots (138 treated, 1,623 untreated). The 1:1 matched sample is based on 5,855 seedlings on 272 plots (136 pairs of treated/untreated plots).

**Supplementary Figure S7:**
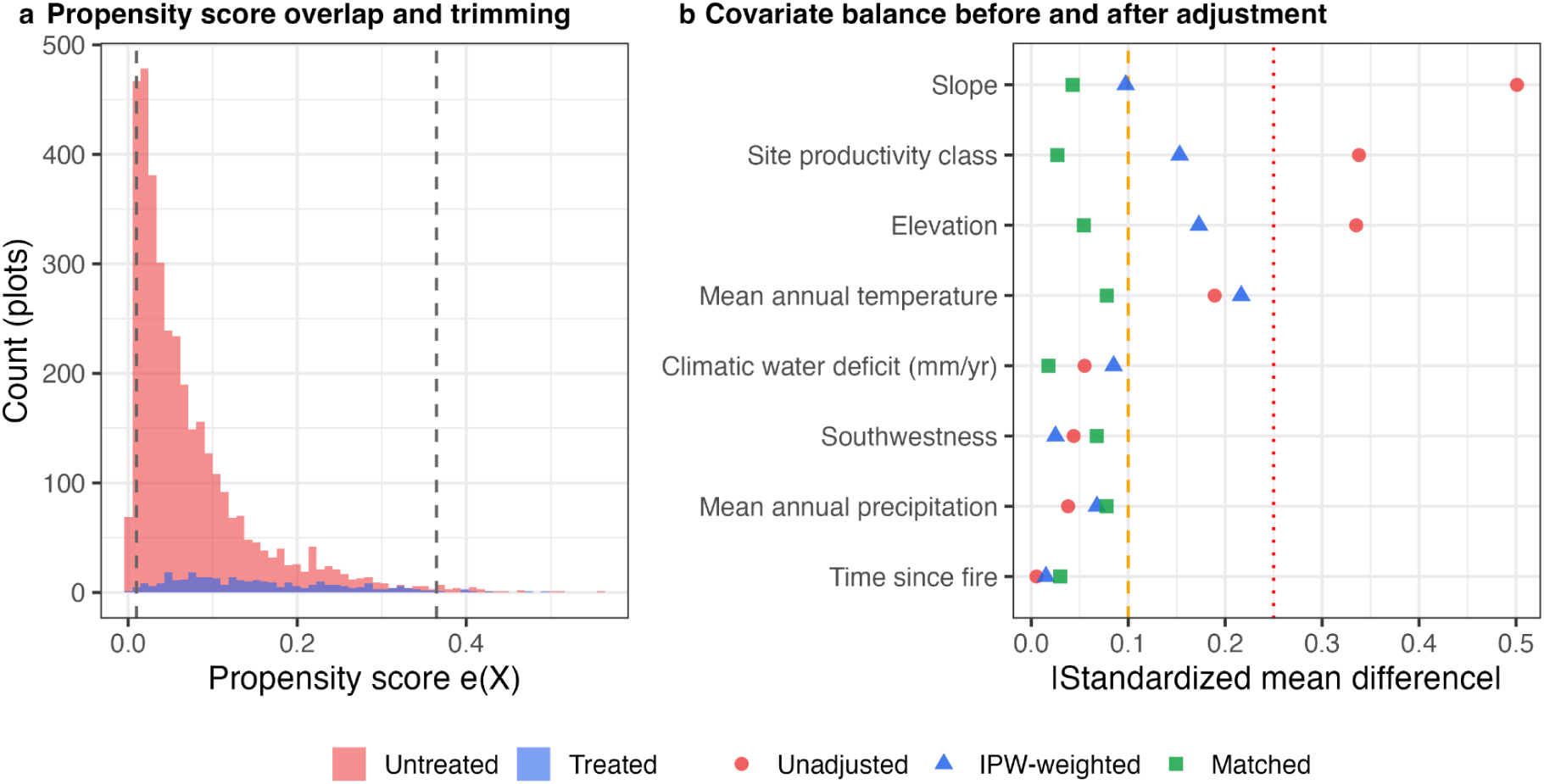
Balancing and diagnostics for the tree dataset. (a) Distribution of the estimated propensity score for untreated (red) and treated (blue) plots. The two vertical dashed lines mark the applied positivity-trimming bounds, where the lower bound lies at e(X) = 0.01 and the upper bound at the 99th percentile of the fitted scores (e(X) = 0.36). Plots outside these bounds are removed (330 of 3,959 plots). The region between the dashed lines is the common support on which every estimate rests. (b) Absolute standardized mean difference between treated and untreated plots for each covariate, unadjusted (red), after stabilized inverse-probability weighting (blue) and after 1:1 nearest-neighbour propensity-score matching with a 0.2 SD caliper (green), computed on the trimmed analysis sample (303 treated, 3,326 untreated plots). SMD represents the treated-minus untreated mean difference divided by the pooled standard deviation. The vertical dashed line at |SMD| = 0.10 is the threshold below which a covariate is considered balanced. The dotted line at 0.25 marks severe imbalance. Four of 8 covariates exceed 0.10 before adjustment, three after inverse-probability weighting and none do after matching. Covariates are ordered by their unadjusted imbalance.

**Supplementary Figure S8:**
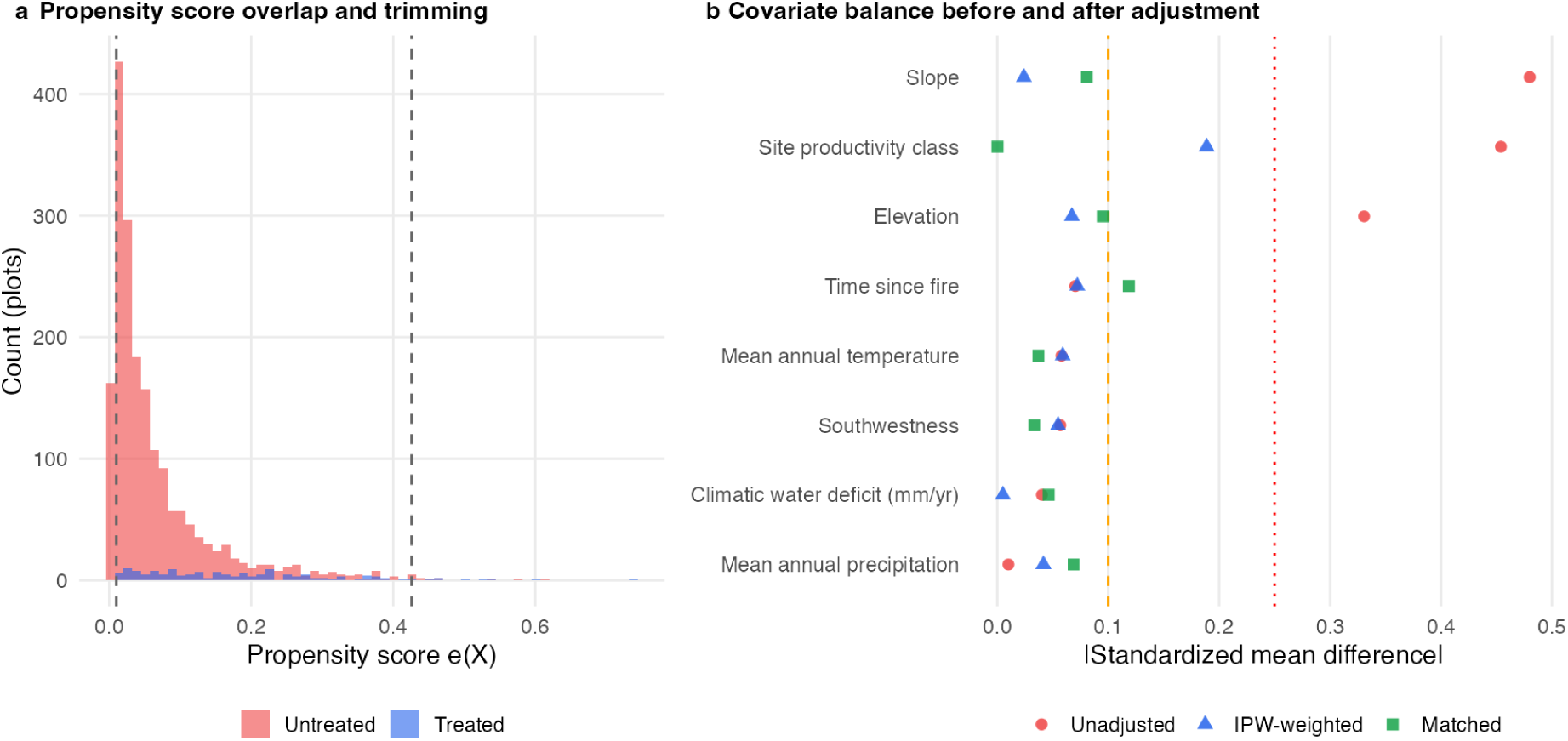
Balancing and diagnostics for the seedling dataset. (a) Distribution of the estimated propensity score for untreated (red) and treated (blue) plots. The two vertical dashed lines mark the applied positivity-trimming bounds, where the lower bound lies at e(X) = 0.01 and the upper bound at the 99th percentile of the fitted scores (e(X) = 0.43). Plots outside these bounds are removed (254 of 2,016 plots). The region between the dashed lines is the common support on which every estimate rests. (b) Absolute standardized mean difference between treated and untreated plots for each covariate, unadjusted (red), after stabilized inverse-probability weighting (blue) and after 1:1 nearest-neighbour propensity-score matching with a 0.2 SD caliper (green), computed on the trimmed analysis sample (138 treated, 1,623 untreated plots). SMD represents the treated-minus untreated mean difference divided by the pooled standard deviation. The vertical dashed line at |SMD|=0.10 is the threshold below which a covariate is treated as balanced. The dotted line at 0.25 marks severe imbalance. Three of 8 covariates exceed 0.10 before adjustment, one after inverse-probability weighting and one after matching (time since fire, which is not a propensity-score covariate). Covariates are ordered by their unadjusted imbalance.

